# The Neovaginal Microbiota, Symptoms, and Local Immune Correlates in Transfeminine Individuals with Penile Inversion Vaginoplasty

**DOI:** 10.1101/2025.03.14.643288

**Authors:** Jorge Rojas-Vargas, Hannah Wilcox, Bern Monari, Pawel Gajer, David Zuanazzi, Ainslie Shouldice, Reeya Parmar, Priscilla Haywood, Vera Tai, Yonah Krakowsky, Emery Potter, Jacques Ravel, Jessica L. Prodger

## Abstract

Transfeminine people (assigned male at birth) often undergo penile inversion vaginoplasty to create vulva, a clitoris and a vaginal canal (referred to as a neovagina). After vaginoplasty, transfeminine people frequently experience gynecological concerns but their etiology is unknown due to a lack of knowledge of the neovaginal microenvironment. We characterized neovaginal microbiota and cytokines in 47 transfeminine participants. Participants self-reported sexual behaviors and symptoms, enabling correlation with bacterial (16S rRNA) and immune profiles. Four distinct clusters of co-occurring bacteria with unique immune profiles were identified. One cluster, which included *Fastidiosipila*, *Ezakiella*, and *Murdochiella*, was abundant, stable, and correlated with lower cytokines. Conversely, another cluster containing *Howardella*, *Parvimonas*, *Fusobacterium*, and *Lawsonella* was linked to higher cytokines. Although *Lactobacillus* was detected, *Lactobacillus*-dominance was rare. These findings underscore the need for evidence-based clinical guidelines tailored to transfeminine gynecologic care, emphasizing the vital role of the neovaginal microbiome in symptom management and sexual health.

## INTRODUCTION

In 2021, more than 100,000 Canadian adults (0.33%) identify as transgender or non-binary, meaning that their gender identity does not align with their sex assigned at birth^1^. Transfeminine people are a group of individuals who were assigned male at birth but do not have a masculine gender identity. There are numerous communities that fall under this umbrella term, including transgender women and non-binary or other gender diverse individuals^2,3^. Transfeminine individuals may pursue gender-affirming medical care, which may include feminizing hormone therapy (e.g., exogenous estradiol and/or progesterone, anti-androgenic drugs) or surgeries^2–9^. Vaginoplasty is a surgery that removes the testis (orchiectomy) and produces a vulva and vaginal canal, referred to as a neovagina in healthcare contexts. The most common technique used is penile inversion vaginoplasty, in which the new vaginal canal is lined with a variable ratio of penile and scrotal skin, dependent on tissue availability and desired depth of the canal^10–15^. Although less common, sigmoid or peritoneal tissue may also be used to augment the length of the vaginal canal^16–18^. Herein, we utilize the term “vagina” to refer to the vaginal canal of those born with a vagina, and we use the term “neovagina” to refer to the surgically created vaginal canal of transfeminine people. Our aim in using two terms to refer to similar organs is to distinguish between vaginal canals lined with epithelia of different origins. The vagina, when estrogenized, has a thick stratified squamous epithelium, the upper layers of which are composed of glycogen-rich cells^19–22^. The penile epithelium is also a stratified squamous epithelium, but is thinner (100-200 vs. 100-300μm) and has a soft-cornified outer layer made up of terminally differentiated keratinocytes in a lipid-rich extracellular matrix (the stratum corneum)^23–25^.

In cisgender males and females, these two epithelia support distinct microbiota. When epithelial cells are shed from the vagina, their glycogen content is metabolized to lactic acid by *Lactobacillus* spp., which lowers the vaginal pH and inhibits colonization by pathogenic organisms, perpetuating *Lactobacillus* dominance^26–28,^. In reproductive aged cisgender females, *Lactobacillus* dominance in the vagina is optimal; however, when the abundance of *Lactobacillus* spp. is low, the vaginal microbiota comprises diverse strict and facultative anaerobes, including *Gardnerella vaginalis, Fannyhessea vaginae* (formerly *Atopobium*), *Mobiluncus* spp., and *Prevotella* spp.^29,30^. In cisgender females, this diverse microbiota is frequently associated with symptoms, including pain, discharge, and malodour, referred to as bacterial vaginosis (BV)^31^. However, even in the absence of symptoms, a diverse microbiota deficient in *Lactobacillus* is associated with altered cytokines and an increased risk of negative sexual and reproductive health outcomes, including sexually transmitted infections, pelvic pain, and obstetric complications^32–34^. Because of the significant health associations of asymptomatic vaginal bacterial dysbiosis, non-optimal vaginal microbiota comprising diverse anaerobes are referred to as “molecular BV”^35,36^.

In contrast, corneocytes of the penile epithelium of cisgender males do not produce glycogen and are packed with keratin bundles and embedded in a lipid matrix, which can be used as a carbon source by bacteria when cells are shed through desquamation^37–40^. Lactobacilli are rare on the penile skin of cisgender males, as are *Gardnerella*, *Fannyhessea*, and *Mobiluncus*^41^. Instead, the sub-preputial space of the uncircumcised penis has a diverse microbiota of Gram-positive and negative anaerobes, including *Prevotella*, *Peptoniphilus*, *Finegoldia*, *Anaerococus*, and *Porphyromonas*^41^. While the bacterial taxa on the penis and in the vagina differ, much like the vagina, a penile microbiota with a high abundance of strict anaerobes is associated with increased local inflammation and risk of adverse sexual health outcomes^41–48^. Circumcision drastically reduces penile anaerobes, promotes colonization with *Corynebacterium*, and concomitantly reduces the risk of negative health outcomes^49–52^.

The impact of circumcision on penile bacteria is evidence of the profound effect that environmental changes can have on the microbiome. Compared to the cisgender male penis, in transfeminine people who have had vaginoplasty the penile skin is invaginated, testosterone levels are decreased, estrogen levels are increased, and direct environmental exposures may be introduced (douching, post-surgical dilation, etc.). Despite the apparent associations between genital bacteria and health outcomes in cisgender people, our knowledge of the neovaginal microbiota of transfeminine people remains limited. Using culture-based methods, a study of transfeminine and non-binary people in Belgium (n=50) found *Bacteroides ureolyticus*, *Corynebacterium* spp., *Enterococcus faecalis*, and *Streptococcus anginosus* to be the most prevalent taxa in the neovagina of Belgian transfeminine people^53^. There have also been two studies that used 16S rRNA gene amplicon sequencing, one in Brazil (n=4, penile skin lined)^54^ and one in the US (n=15, unspecified tissue source)^55^. Both reported the neovagina to comprise diverse bacterial communities with a high prevalence of anaerobes typical of the uncircumcised penis, including *Porphyromonas*, *Peptoniphilus, Prevotella*, *Peptostreptococcus*, and *Finegoldia.* Interestingly, the US-based study also observed high prevalence of *Lawsonella clevelandensis*, which is not commonly observed on the penis of cisgender males. Although *Lactobacillus* can be detected at low relative abundance in most neovaginal samples^56,57^, a common observation from these studies is that *Lactobacillus* dominance post-penile inversion vaginoplasty is rare.

Gynecological symptoms are commonly reported by transfeminine people with vaginoplasty, including odour, discharge, and itching – all indicators of BV in cisgender females^27,58,59^. However, when methods to diagnose BV (*e.g.*, Nugent score) are applied to the neovagina they return inconclusive results, and antibacterial treatments used to treat BV in cisgender females (*e.g.*, metronidazole) are often ineffective for transfeminine individuals. Understanding the neovaginal microenvironment is crucial to developing effective diagnostics and treatments for transfeminine people. This study characterized the composition and structure of the neovaginal microbiota of 47 transfeminine participants and examined associations with neovaginal inflammation, symptoms, and exposures.

## RESULTS

### Participant demographics

This study included 47 transfeminine individuals (TF), 35 reproductive age cis female individuals (rCF), and 56 cis male individuals (CM). Of the 47 TF participants, 45 provided swab samples at all three weekly timepoints and two provided samples at only 2 timepoints (total n=139 TF samples). 16S rRNA gene analysis was successfully completed 133/139 TF samples, pH reported for 127 samples, and immune analysis was successful in 133 samples. Participant characteristics are listed in **Supplemental Table 1**. TF participants were a mean age of 41 years (range 23-69) and predominantly identified as “white” (89%). Participants had undergone vaginoplasty, on average, 4.3 years prior to enrollment (range 1-19 years) and approximately half of participants were circumcised prior to vaginoplasty (51.1%). All TF participants reported using feminizing hormone therapy at all study visits. TF participants were significantly older and more likely to identify as white compared to CM and rCF participants.

### Microbiota of the penile-skin lined neovagina

Initial taxonomic assignment was made to a maximum depth of genus and 16S rRNA gene amplicon sequence variants (ASV) were collapsed into a total of 240 taxa, representing 182 genera and 38 taxa that could only be classified at the level of family or higher. The median number of genera detected per sample was 36 and the range was 8-64. The core neovaginal microbiota (**Figure 1**), defined as genera present in 50% of samples with at least 1% relative abundance, was found to be comprised of 11 genera: *Peptoniphilus* (98.5% prevalence, 10.68% median relative abundance), *Prevotella* (97.7% prevalence, 7.78% median relative abundance), *Hoylesella* (97.0% prevalence, 7.77% median relative abundance), *Porphyromonas* (92.5% prevalence, 6.76% median relative abundance), *Anaerococcus* (97.7% prevalence, 3.04% median relative abundance), *Dialister* (95.5% prevalence, 2.36% median relative abundance), *Fenollaria* (78.9% prevalence, 1.86% median relative abundance), *Ezakiella* (84.21% prevalence, 1.35% median relative abundance), *Varibaculum* (91% prevalence, 1.19% median relative abundance), *Campylobacter* (92.5% prevalence, 1.13% median relative abundance), and *Fusobacterium* (76.7% prevalence, 1.01% median relative abundance).

**Figure 1.**
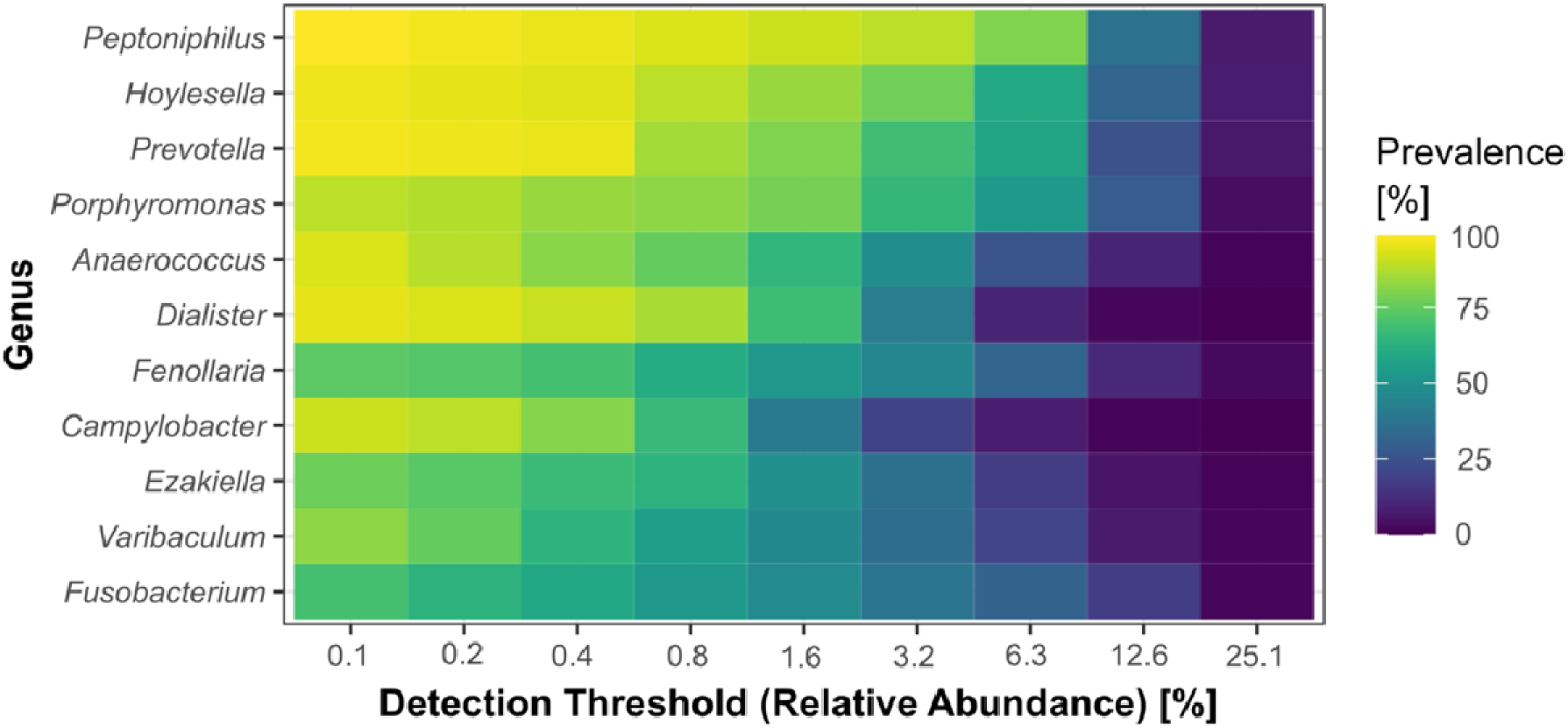
The core microbiota of the neovagina of transfeminine people with penile inversion vaginoplasty. Bacteria were characterized in 133 neovaginal swabs from 47 participants using 16S rRNA gene amplicon sequence analysis. The core microbiota was defined as genera present in 50% of samples with at least 1% relative abundance (created using the core_members function, microbiome R package).

Within the 182 genera, further species-level assignments could be made for 91 genera (**Supplemental Table 2**). The most common species of *Peptoniphilus* was *P. harei* (96.2% prevalence and 1.73% median relative abundance); of *Hoylesella* was *H. timonensis* (previously in the genus *Prevotella*; 88.72% prevalence and 5.91% median relative abundance); and of *Prevotella* was *P. disiens* (50.38% prevalence and 0.007% median relative abundance).

### The neovaginal microbiota is distinct from the penile and vaginal microbiota

To contextualize the neovaginal microbiota, we also analyzed vaginal swabs from rCF and sub-preputial swabs from uncircumcised CM. Comparisons between the three anatomical sites were made at a maximum taxonomic depth of genus. Compared to the rCF vagina, the neovagina had a similar median number of genera (36 vs. 38 genera in TF vs. rCF, p=0.346), but significantly higher bacterial diversity (Shannon Diversity Index: 2.55 vs. 0.71 in TF vs. rCF, p<0.001). Compared to the CM sub-preputial space, neovaginal samples had significantly fewer genera (median 36 vs. 41 genera in CM, p<0.0001), but significantly higher diversity (Shannon Diversity Index 2.55 vs. 2.25 in CM, p<0.001). **Figure 2** and **Table 2** compare the bacterial composition of the microbiota at these three anatomical locations.

**Figure 2.**
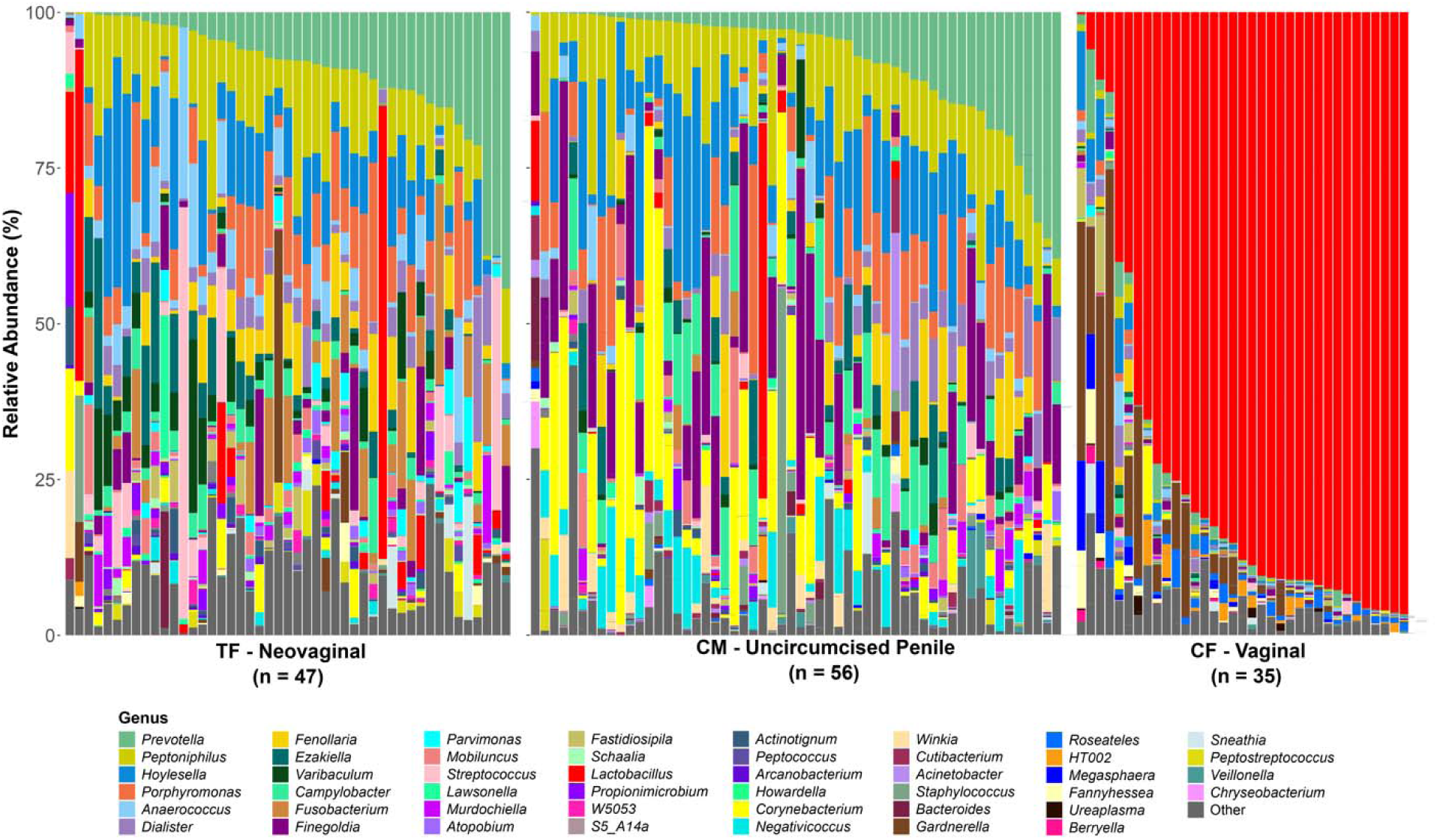
Relative abundances of bacteria in the TF neovagina, uncircumcised CM penis, and reproductive-aged rCF vagina. Bacteria in genital swabs from transfeminine (TF, n=47), cis male (CM, n=56), and cis female (rCF, n=35) participants were characterized by 16S rRNA gene amplicon sequencing. The relative abundances of the top 30 most abundant genera in each anatomical location (by median relative abundance) are displayed as stacked bar plots (total 46 genera). Each vertical bar represents one participant; for TF participants who contributed multiple samples, only the first available timepoint is presented. A bar plot with data from all TF participant study visits (n=133) is presented in **Supplemental Figure 1**.

**Table 2.**
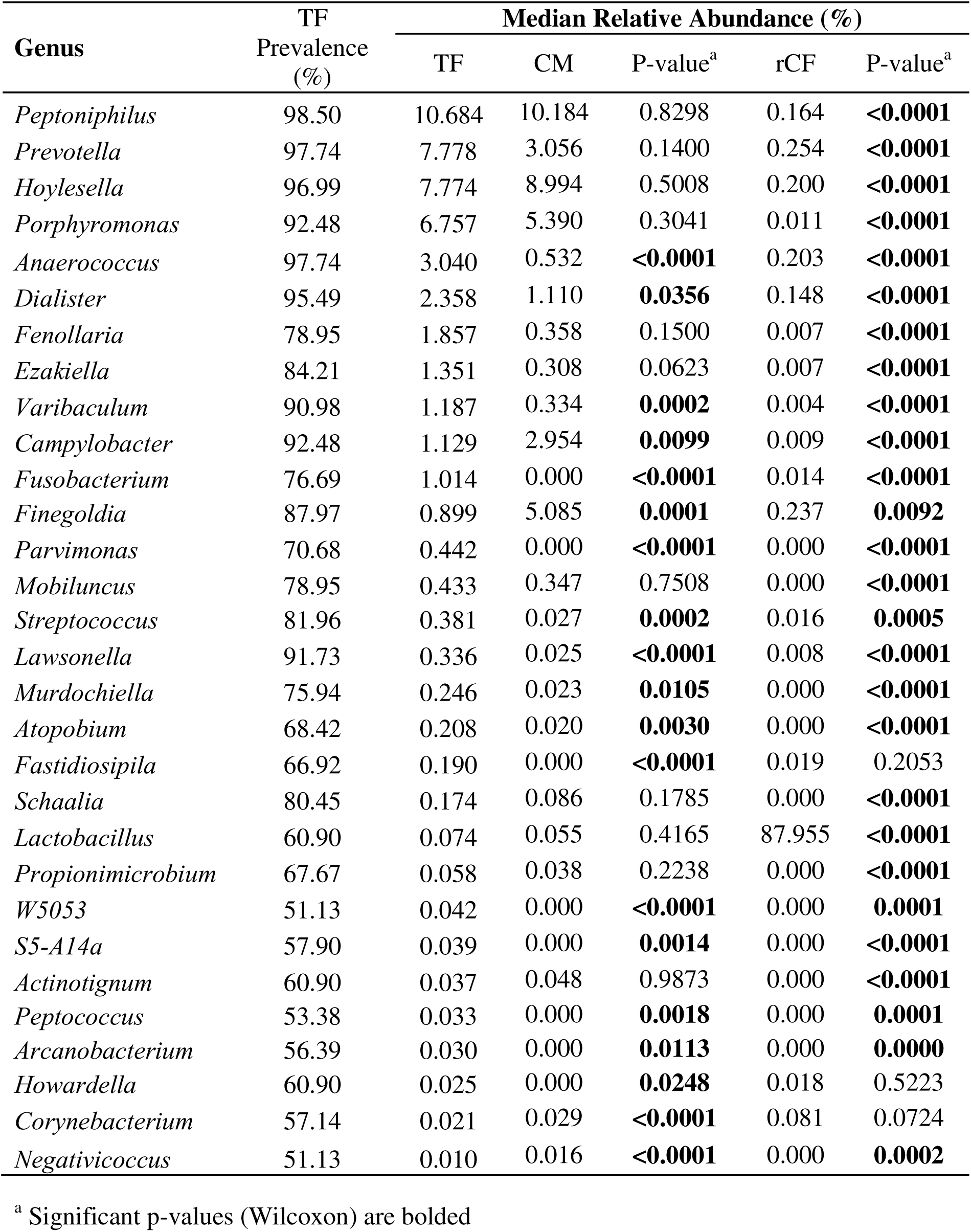
Thirty most abundant bacterial genera in the neovagina of transfeminine (TF) participants, compared to the penile and vaginal microbiota of cis male (CM) and reproductive age cis female (rCF) participants.

As previously reported in other cohorts, the majority of rCF had microbiota dominated by *Lactobacillus*, with a minority of rCF having reduced lactobacilli and instead a diverse microbiota consistent with molecular BV, including *Gardnerella*, *Hoylesella* (species previously belonging to *Prevotella*), *Megasphaera*, and *Prevotella.* The most prevalent and abundant taxa on the uncircumcised penis of CM were *Peptoniphilus*, *Hoylesella* (species previously belonging to *Prevotella*), *Porphyromonas, Finegoldia,* and *Prevotella*. However, a continuum exists, where men with increasing *Corynebacterium* also had increasing *Staphylococcus, Negativicoccus, Anaerococcus*, and *Finegoldia*, with decreasing *Porphyromonas, Hoylesella, Peptoniphilus*, and *Prevotella*.

The four most abundant genera in the neovagina – *Peptoniphilus*, *Prevotella, Hoylesella,* and *Porphyromonas* – were not significantly different in median relative abundance from CM penile samples (**Table 2**). Of the 30 most abundant genera in the neovagina, 19/30 were significantly different in median relative abundance between the neovagina and penis, while 27/30 were significantly different between the neovagina and vagina. *Lactobacillus* was frequently detected in neovaginal samples (82/133 samples, 60.9%), but at a median relative abundance that was substantially lower than the rCF vagina (0.07 vs. 87.96% median relative abundance, p<0.001) and more like the median relative abundance on the uncircumcised CM penis (0.06%, p=0.4).

### Associations between neovaginal bacteria and immunology

To gain insight into the relevance of neovaginal microbiota composition to TF health, we examined associations between neovaginal bacteria and local inflammation. As local immune responses are expected to vary temporally with changes in the neovaginal microbiota, data from all timepoints are included in subsequent analyses (controlling for longitudinal sampling). Concentrations of immune analytes in neovaginal swab eluants are presented in **Supplemental Table 3**. Of the 19 immune analytes assayed, 7 cytokines (IL-1α, IL-1β, IL-6, IL-8, MIG, MIP-1β and RANTES) were quantifiable in at least 40% of samples and were included in subsequent analyses.

To identify microbiota that may influence neovaginal immune responses, we generated a correlation matrix (**Figure 3**) of the 30 most abundant genera in the neovagina, examining correlations (i) between relative abundances of bacterial genera and (ii) between relative abundance of bacterial genera and concentration of immune analytes.

**Figure 3.**
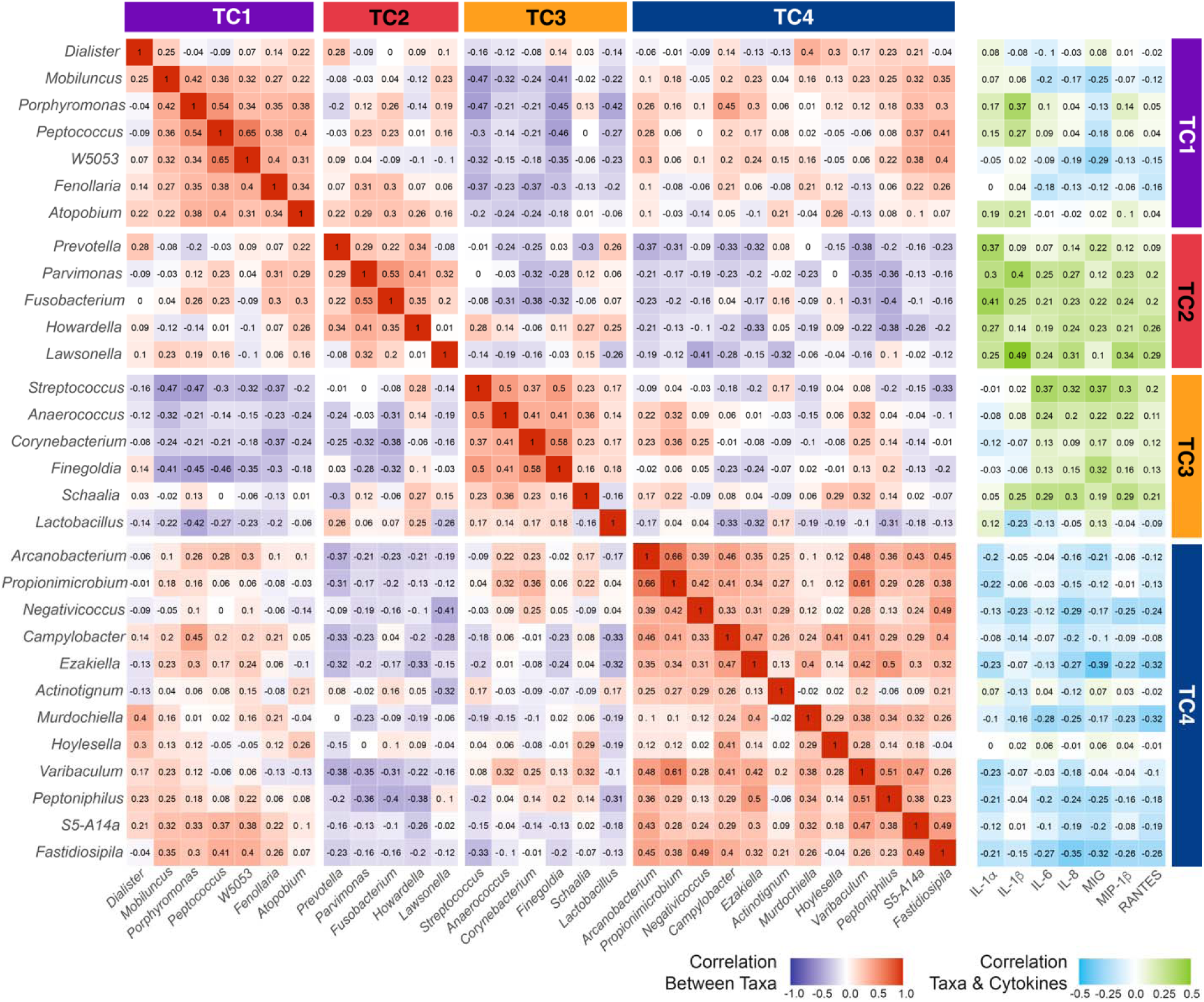
Bacterial genera in the neovagina co-correlate in four distinct clusters. Neovaginal bacteria (133 samples representing 47 participants) were characterized by 16S rRNA gene amplicon sequencing and cytokines were quantified by multiplex immunoassay (Luminex). Spearman’s correlation coefficients between the 30 most abundant neovaginal genera (median relative abundance) are displayed in a correlation matrix. K-means clustering identified four Taxa Clusters (TC). Spearman correlation coefficients between each genus and each cytokine are presented in the matrix on the right. P-values associated with Spearman’s correlations are presented in **Supplemental Tables 4 and 5**.

K-means clustering of Spearman correlations between genera identified 4 unique clusters of co-occurrent bacteria (referred to as Taxa Clusters, or TC), with genera in TC2 and 3 generally positively associated with cytokines, those in TC4 generally negatively associated with cytokines, and genera in TC1 showing mixed associations (p-values for Spearman correlations in **Supplemental Tables 4 and 5**).

TC2 is comprised of five genera – *Prevotella, Parvimonas*, *Fusobacterium, Howardella*, and *Lawsonella –* all of which were significantly positively correlated with IL-1α and at least one other cytokine. In addition to IL-1α, *Prevotella* abundance was positively associated with only MIG. Other genera in TC2 were associated more broadly with pro-inflammatory markers (IL-6, IL-8, MIG, MIP-1β, RANTES): *Parvimonas* was significantly associated with 3/5 of these cytokines and *Fusobacterium, Howardella*, and *Lawsonella* were significantly associated with 4/5. Genera in TC2 were generally significantly negatively correlated with taxa in TC4, except for *Hoylesella,* which showed no significant correlations with TC2 taxa.

TC3 comprises six genera, of which four – *Schaalia* (formerly *Actinomyces* spp.), *Streptococcus, Anaerococcus,* and *Finegoldia –* were significantly positively correlated with cytokines. Of these four, *Schaalia*, *Streptococcus,* and *Anaerococcus* were positively associated with multiple pro-inflammatory markers, while *Finegoldia* was correlated significantly with only MIG. TC3 also included *Corynebacterium,* which showed no significant correlations with cytokines, and *Lactobacillus*, which was negatively correlated with IL-1β. The abundances of most TC3 bacteria were significantly negatively correlated with the abundances of TC1 bacteria, except for *Schaalia,* which showed no significant correlations with TC3 bacteria.

TC4 is the largest TC, comprised of 12 genera, of which eight were significantly negatively correlated with at least one cytokine. The most striking negative associations with pro-inflammatory markers were observed with *Fastidiosipila* (significant negative correlations with 5/5 pro-inflammatory markers: IL-6, IL-8, MIG, MIP-1β, RANTES), *Ezakiella* (significant negative correlations with 4/5), and *Murdochiella* (significant negative correlations with 4/5). No significant positive associations were observed between any TC4 bacteria and any cytokines.

TC1 comprises seven genera that tended to correlate negatively with IL-1α and IL-1β, but positively with other cytokines, although few correlations were significant. *Porphyromonas* and *Peptococcus* were significantly negatively associated with IL-1β and *Mobiluncus* and *W5053* were significantly positively associated with MIG.

For each swab, the relative abundances of the genera comprising each TC were summed to obtain a relative abundance for the TC as a whole. Overall, TC4 bacteria were the most abundant, with a median relative abundance of 32.80%, followed by TC1 (16.95%) and TC2 (13.77%), while TC3 bacteria were the least abundant (9.01%). Hierarchical clustering of participant samples based on TC relative abundances grouped participants into 6 clusters, denoted by α, β, χ, δ, ε, and φ in **Figure 4**. Individuals belonging to clusters α and β – characterized by high levels of TC1 and TC4, respectively – had lower levels of cytokines, while individuals in clusters χ, δ, and ε had comparatively higher cytokines (**Figure 4C**). Of note, cluster φ had the lowest median concentration of all cytokines; most individuals in cluster φ had high abundance of *Lactobacillus*. To examine the week-to-week stability of neovaginal bacterial communities, we calculated the probability of a participant’s sample transitioning from one cluster to another between two weekly samples (Markov probabilities, **Figure 4D**). Stability was high for individuals in clusters α, β, χ, δ, and ε, with >64% of participants remaining in the same cluster between two weeks. However, individuals in cluster φ displayed less stability, with a probability of 33% of remaining in cluster φ and 50% of transitioning to cluster α.

**Figure 4.**
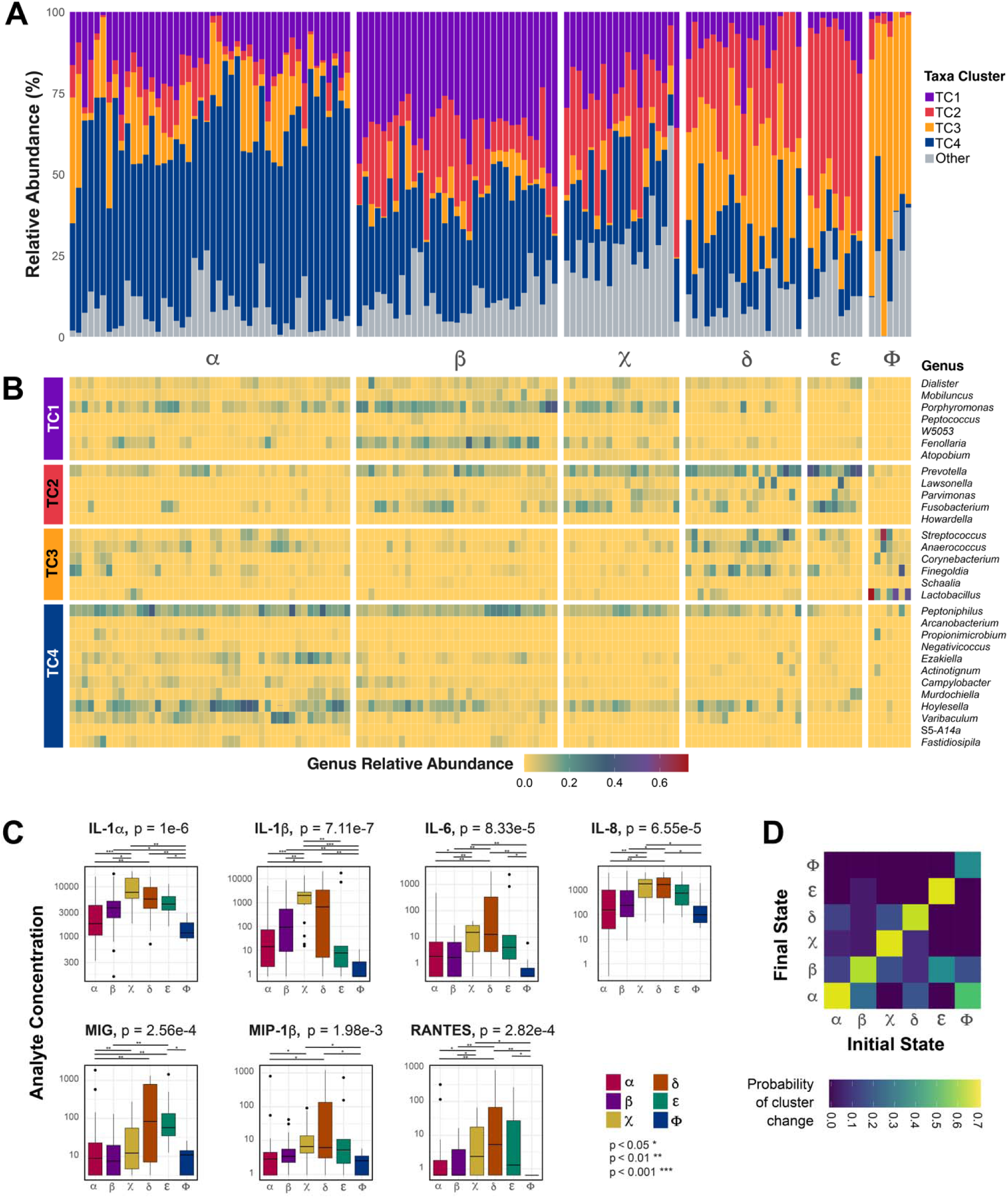
Relative abundance of Taxa Clusters varies substantially between transfeminine individuals. Relative abundances of the genera comprising each of the four Taxa Clusters (TC) were summed to generate an overall relative abundance for each TC. Samples (n=133 from 47 participants) were grouped using hierarchical clustering (denoted by α, β, χ, δ, ε, φ; clustering by hc and cutree) based on their relative abundance of each TC **(A)**. Genera comprising each sample are displayed in **B**. Median cytokine concentration of participants in each cluster (α, β, χ, δ, ε, φ) are displayed in **C**; Kruskal-Wallis test was used to test for differences in distribution, followed by Dunn’s test for pairwise comparisons. The Markov probability that a participant’s sample belongs to a particular cluster (α, β, χ, δ, ε, φ), given the cluster they belonged to in the previous week, is displayed as a heatmap matrix in **D**.

### Correlation between neovaginal bacteria and participant characteristics

To explore potential determinants of neovaginal bacteria, we examined associations between the median relative abundance of each TC with participant’s age, time since vaginoplasty, pre-vaginoplasty circumcision status, and neovaginal behavioural profile (**Supplemental Table 6**) using zero-inflated beta regression models and accounting for longitudinal sampling. No significant associations were observed between the relative abundance of any of the TCs and age or time since vaginoplasty (all p > 0.3, **Supplemental Table 7**). However, pre-vaginoplasty circumcision status was significantly associated with neovaginal bacterial composition. Participants who were circumcised prior to vaginoplasty had significantly less TC1 bacteria (median relative abundance 14.53 vs. 21.96 %, p < 0.001) and TC2 (9.98 vs. 15.83 %, p < 0.001) and significantly more TC3 (10.99 vs. 5.93 %, p < 0.001), and TC4 (39.50 vs. 26.93 %, p < 0.001) bacteria (**Figure 5A**, **Supplemental Table 7**). Specifically, prior circumcision was associated with lower (0.05 < p < 0.25, FDR adj.) *Prevotella*, *Parvimonas*, and *Moryella* (previous *Prevotella* spp.), and higher abundances of *Corynebacterium*, *Finegoldia*, *Varibaculum*, *Propionimicrobium*, *Facklamia*, and *Gleimia* (**Figure 5B**). Species-level assignments revealed participants who were circumcised prior to vaginoplasty had significantly higher abundances of *P. harei* and *Fastidiosipila sanguinis* (**Figure 5B**; FDR adj. p<0.05).

**Figure 5.**
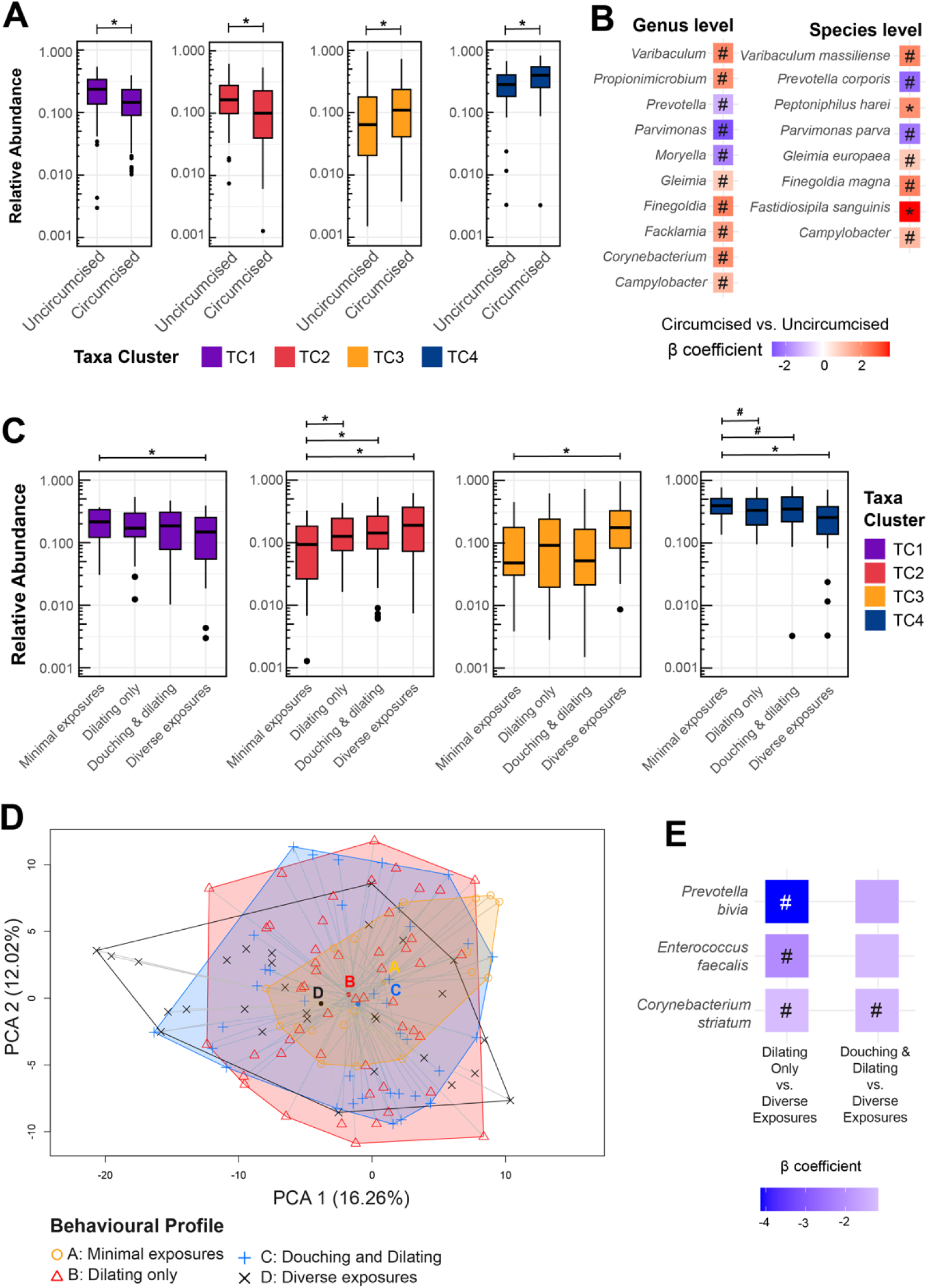
Neovaginal bacteria differ based on pre-vaginoplasty circumcision status and neovaginal behavioural profile. Relative abundances of the genera belonging to each taxa cluster (TC) were summed (n=133 samples from 47 individuals). **(A)** Median relative abundance and interquartile range of TC relative abundance, stratified by pre-vaginoplasty circumcision status. Asterisks (*) represent significant differences calculated using beta regression model, p < 0.05. **(B)** Differential abundances of genera and species based on pre-vaginoplasty circumcision status were calculated with MaAsLin2. Taxa identified as being significantly different are displayed (beta regression model with false discovery rate adjustment; asterisk (*) denotes adj. p<0.05 and crosshatch (#) denotes 0.05 < adj. p < 0.25). **(C)** Median TC relative, stratified by behavioural profile. Asterisks denote significant differences calculated using beta regression model (p < 0.05), crosshatches denote 0.05 < p < 0.25. **(D)** Principal Component Analysis (PCA) analysis of 16S rRNA gene amplicon sequence read counts with a centered log-ratio (CLR). Within-group dispersion (Aitchison distances) is depicted by shading. **(E)** Differential abundances of genera and species between behavioural profiles were identified with MaAsLin2 and taxa identified as being significantly different are displayed.

Neovaginal behavioural profiles were previously defined in this cohort using hierarchical clustering, which revealed participants fall into four distinct clusters based on their self-reported neovaginal practices/exposures: (1) limited neovaginal exposures, (2) regular neovaginal dilating, (3) regular neovaginal dilating and douching, and (4) diverse exposures^60^. Participants who reported diverse exposures had significantly higher abundances of TC2 and TC3 bacteria, and significantly lower abundances of TC1 and TC4 bacteria, compared to participants who reported limited neovaginal exposures (**Figure 5C**). Participants who dilated only (no douching), as well as participants who dilated and douched, had significantly (p<0.05) higher abundance of TC2 bacteria compared to participants with limited exposures and tended to have higher abundance of TC4 bacteria (p=0.05) compared to participants with diverse neovaginal exposures (medians, beta coefficients, and p-values in **Supplemental Table 9**).

To explore the association between behavioural profiles and microbiota dispersion and composition, we performed a Principal Component Analysis (PCA) analysis on the 16S rRNA gene amplicon sequence read counts with a centered log-ratio (CLR) transformation (**Figure 5D**). While global PERMANOVA revealed significant differences in dispersion between behavioural profiles (p=0.023), post-hoc pairwise comparisons were not significant after adjusting for false discovery rate (adj. p<0.05). Community structure also differed significantly between behavioural profiles (global PERMANOVA p = 0.001, r^2^=0.043). Post-hoc differential abundance analysis revealed participants with diverse exposures tended to have higher relative abundances of *P. bivia* (β = -4.13, adj. p = 0.15), *E. faecalis* (β = -2.07, adj. p = 0.21), and *Corynebacterium striatum* (β = -1.17, adj. p = 0.15, **Figure 5E**) than participants who douched and dilated.

### The neovaginal microbiota is associated with patient symptoms

To explore relationships between neovaginal bacteria and symptoms, we examined neovaginal pH at the time of sample collection and self-reported bleeding, discharge, itching/burning, malodour, and pain in the 7 days prior to sample collection. Neovaginal pH values ranged from 3 – 8; the most commonly reported pH was 5.5, with 41.7% of samples having this value (53/127; **Supplemental Figure 2**). After controlling for longitudinal samples from the same individual, the relative abundance of TC4 bacteria was positively associated with neovaginal pH (p=0.026, **Supplemental Table 7)**. No significant associations were found between pH and other TC (all p > 0.05). Symptoms were reported at 26.3% of sample collections; the most common symptom was neovaginal malodour (defined as an odour that was bothersome or unpleasant), reported for 19/133 samples, followed by bleeding (n=10), discharge (n=7), and itching or burning (n=6). Pain (n=2) was less commonly reported (**Supplemental Table 8**).

The strongest association between neovaginal bacteria and symptoms was observed for malodour. Compared to asymptomatic study visits, malodour was associated with significantly lower abundance of TC2 bacteria (6.35 vs. 13.49% median relative abundance, p= 0.005) and significantly higher abundance of TC4 bacteria (50.67 vs. 32.29%, p<0.001, **Figure 6A** and **Supplemental Tables 6 and 7**).

**Figure 6:**
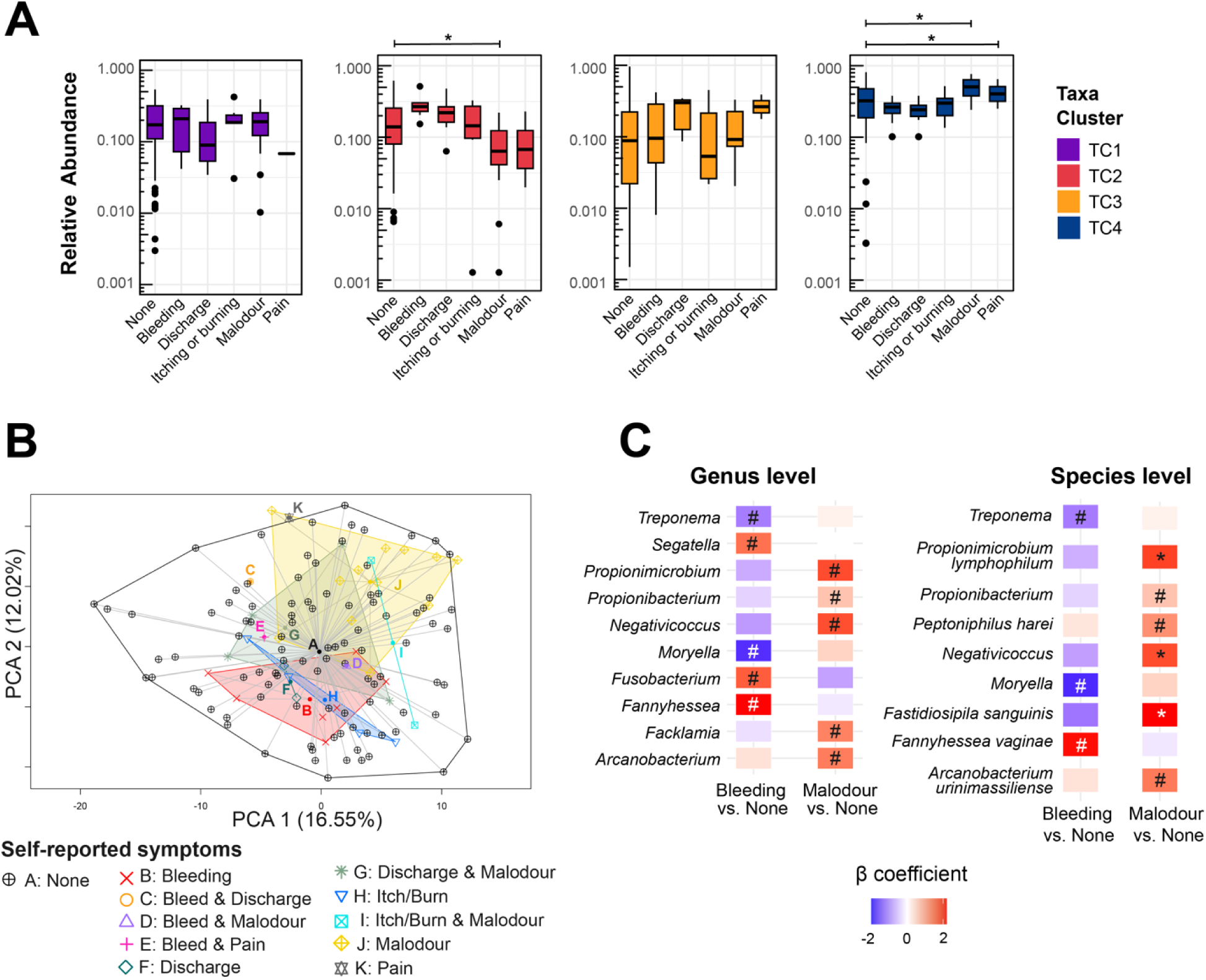
Neovaginal malodour is associated with bacteria. Participants (n=47) self-reported symptoms they experienced in the week prior to neovaginal swab collection (n=133 samples). Relative abundances of the genera belonging to each taxa cluster (TC) were summed and the median TC relative abundance stratified by symptom type is presented in **A** (n=133 samples from n=47 participants). Asterisks (*) denote significant differences calculated using beta regression model (p < 0.05), crosshatches (#) denote 0.05 < p < 0.25. Principal Component Analysis (PCA) analysis of 16S rRNA bacterial counts with a centered log-ratio (CLR) transformation is displayed in **B**. Within-group dispersion (Aitchison distances) is depicted by shading. Differential abundances of genera and species between asymptomatic and symptomatic study visits were calculated with MaAsLin2. Taxa identified as being significantly different are displayed in **C**.

We performed a PCA analysis to explore the association between self-reported symptoms and microbiota dispersion and composition (**Figure 6B**). Study visits where participants reported symptoms fell within the dispersion of asymptomatic visits. However, a global PERMANOVA was significant (p=0.001, r^2^=0.483) and post-hoc pairwise comparisons revealed that study visits where only malodour was reported were significantly more tightly clustered than asymptomatic visits (adj. p=0.021). Significant differences in dispersion were also observed between study visits with only discharge vs. those with itching/burning and malodour (adj. p=0.021), but observation sizes in these two groups were very small (n=2 in each group). Study visits where bleeding was reported tended to be more tightly clustered than asymptomatic visits (adj. p=0.091).

We next examined differences in microbiota composition across study visits (beta diversity, global PERMANOVA p = 0.006). Post-hoc pairwise comparisons revealed that study visits where only malodour was reported had a significantly different microbiota composition compared to asymptomatic visits (adj. p=0.020), visits with only itching/burning (adj. p=0.020), and visits with only bleeding (adj. p=0.005). After controlling for repeat sampling, differential abundance analysis identified five genera that tended to be more abundant (adj. p<0.25) in study visits with malodour compared to asymptomatic visits: *Propionimicrobium*, *Propionibacterium*, *Negativicoccus*, *Facklamia*, and *Arcanobacterium* (**Figure 6C**). Species-level taxonomic assignments identified *Propionimicrobium lymphophilum* and *F. sanguinis* as significantly associated with malodour (adj. p<0.05). Compared to asymptomatic study visits, visits with bleeding tended to have a higher abundance of *Segatella*, *Fusobacterium* and *Fannyhessea*, and a lower abundance of *Treponema* and *Moryella* (adj. p<0.25; *Moryella,* previously *Prevotella* spp.). Species level taxonomic assignments identified *F. vaginae* (formerly *A. vaginae*) as being enriched in samples with bleeding (adj. p<0.25).

## DISCUSSION

Our findings demonstrate that neovaginal microbiota of TF after penile inversion vaginoplasty differ substantially from vaginal communities of rCF, and instead more closely resembled those found in the sub-preputial space of the uncircumcised penis, with some key differences. This suggests that the tissue used to create the neovagina strongly influences microbiota composition. We identified four clusters of co-occurrent neovaginal bacteria (referred to here as taxa clusters, TC) that were associated with distinct immune profiles. These observations suggest interplay between bacteria and the neovaginal tissue, consistent with observations in the rCF vagina and the penile sub-preputial space^41^.

The beneficial effects of *Lactobacillus* dominance in the rCF vagina are well established^19,61,62^. While *Lactobacillus* was detected in over half of TF neovaginal samples, it was generally detected at low abundance and *Lactobacillus* dominance was rare. Despite the rarity, there is some evidence to suggest that high *Lactobacillus* abundance might have beneficial effects on the neovaginal epithelium. In the cohort as a whole, the relative abundance of *Lactobacillus* correlated with decreased levels of IL-1β but did not correlate significantly with other pro-inflammatory cytokines (IL-6, IL-8, MIG, MIP-1β, RANTES). However, when clustered based on TC relative abundance, individuals with high *Lactobacillus* abundance clustered together, and this cluster (φ) had the lowest median concentration for all cytokines assessed. While his suggests a potential beneficial role for *Lactobacillus*, this community type was both rare and had low week-to-week stability, suggesting attempts to promote *Lactobacillus* dominance may face difficulties.

In contrast, we identified an assemblage of co-occurrent bacteria in the neovagina that were highly abundant, associated with decreased markers of inflammation, and displayed high week-to-week stability. Of the four TC identified, TC4 bacteria were the most abundant, making up 38% of all bacterial sequences. TC4 also had a favourable immune profile: abundances of TC genera inversely correlated with concentrations of pro-inflammatory cytokines, with the strongest immunomodulatory associations observed for *Fastidiosipila*, *Ezakiella*, and *Murdochiella*. Some TF had near complete TC4 dominance, characterized by a uniquely high abundance of *Hoylesella* (predominantly *H. timonensis*, previously in *Prevotella*), *Ezakiella,* and *Varibaculum*. Participants with this type of microbiota had low levels of all cytokines measured. Additionally, abundance of TC4 bacteria was negatively correlated with TC2 and TC3 bacteria, which are associated with immune activation. It is impossible to discern from this observational study if TC4 bacteria directly dampen neovaginal inflammation or if their negative association with cytokines is due to their inverse relationship with inflammatory bacteria. However, both actions – exclusion of inflammatory bacteria or direct immunomodulatory effects – are characteristics of an immunologically optimal neovagina microbiota. Further *in vitro* studies should explore causal relationships between TF bacteria and inflammation, and antagonism between species.

In keeping with observations in cisgender males (CM), abundances of *Prevotella* and *Howardella* were positively associated with neovaginal cytokines. In the neovagina, *Prevotella* and *Howardella* co-occur with *Parvimonas*, *Fusobacterium*, and *Lawsonella* (TC2), and this group of bacteria was most strongly and broadly associated with markers of inflammation; all TC2 genera were positively correlated with multiple cytokines. Species belonging to this group of genera have been linked to mucosal inflammation at other sites, including the rCF vagina (*Prevotella, Fusobacterium* and *Parvimonas*)^63–66^ and the oral cavity (specifically *Parvimonas micra* and *Fusobacterium nucleatum,* both found in the neovagina)^50,67,68^, and several case studies have implicated *L. clevelandensis* (detected in >90% of neovaginal samples) in serious internal infection and inflammation^69,70^. These data suggest that TC2 bacteria are non-optimal in the neovagina.

In CM, an optimal sub-preputial microbiota has lower abundances of *Prevotella*, *Dialister*, *Peptostreptococcus*, and *Howardella,* and higher abundances of *Corynebacterium*, *Finegoldia*, *Anerococcus*, and *Negativicoccus*^41^. CM with this community type have low sub-preputial IL-8, low density of foreskin T cells, and a reduced risk of acquiring HIV-1^41^. There are similarities between this optimal penile community and neovaginal TC3, which contains *Corynebacterium*, *Finegoldia*, and *Anaerococcus,* although TC3 also contains *Lactobacillus*, *Streptococcus,* and *Schaalia* (previously *Actinomyces* spp.). While *Corynebacterium* and *Finegoldia* were not strongly associated with increased cytokines themselves, they co-occur with *Streptococcus* and *Anaerococcus*, which were associated with increased neovaginal cytokines (IL-6, IL-8, MIG, and MIP-1β, but not IL-1α/β). *Schaalia,* also clustering in TC3, was present at low abundance but was strongly associated with increased IL-1β, IL6, IL-8, MIG, and MIP-1β. *Schaalia* spp. are common at mucosal sites (rCF vagina, oropharynx, gastrointestinal tract), but can cause serious invasive infection (actinomycosis) in the setting of epithelial injury^71^. The observation that optimal penile bacteria co-occur with inflammation-associated genera in the neovagina suggests that, despite some similarities, microbiota in the neovagina and sub-preputial space are distinct.

Interestingly, BV in the rCF vagina is not associated with ubiquitously increased cytokines, but instead shows a distinctive pattern of increased IL-1α/β, marginally increased or unchanged IL-6 and IL-8, and decreased IP-10^64,66,72,73^. Resolution of BV following metronidazole treatment of rCF in sub-Saharan Africa leads to decreased IL-1α/β, but increased IP-10, MIP-1α, and MIG (associated with *Lactobacillus iners* colonization)^74–77^. The neovaginal immune profile observed with TC1 bacteria – *Dialister*, *Mobiluncus*, *Porphyromonas*, *Peptococcus*, *W5053*, *Fenollaria* and *Atopobium* – is similar to BV in CF, with high IL-1α/β but decreased chemokines. While *Dialister* is commonly associated with BV, the other TC1 genera are not typically associated with BV. Of note, *Atobopium* in TC1 does not include *A. vaginae*, frequently associated with BV, as this species was recently re-classified as *F. vaginae*. *Atopobium* in TC1 is predominantly *A. deltae*, with low abundance of *A. minutum* and other uncharacterized *Atopobium* spp. Despite differences in bacteria, the unique cytokine pattern suggests that TC2 bacteria may induce a neovaginal immune response that is similar to that induced by BV in the rCF vagina.

Despite clear associations between neovaginal bacteria and cytokines, few significant associations were observed between bacteria and symptoms. The strongest associations observed were with neovaginal malodour, which was associated with high abundances of *P. lymphophilum*, *F. sanguinis*, and *Negativicoccus* spp. – all belonging to TC4 and associated with reduced inflammation. To our knowledge, these species have not been previously associated with vaginal malodour, halitosis, or skin malodour. Additional research on neovaginal metabolites is needed to better understand the drivers of transfeminine neovaginal malodour. Of note, neovaginal malodour was the most commonly reported symptom, providing the most power to detect associated bacteria. Studies specifically targeting TF experiencing symptoms may be necessary to determine if other BV-like symptoms are associated with neovaginal bacteria.

While this study’s short follow-up and observational nature prevent causal inference, some hypothesis-generating observations can be made on factors that may influence the neovaginal microbiota. Pre-vaginoplasty circumcision status was associated with neovaginal bacteria, and it is tempting to speculate that the penile microbiota may seed and thus influence neovaginal bacteria. However, TF who were circumcised would also have had less penile skin available, necessitating the use of a higher proportion of scrotal skin to provide depth to the neovaginal canal. Scrotal skin is physiologically different from penile / foreskin tissue, notably in the presence of hair follicles, which may influence colonizing bacteria^78^. We also observed associations between neovaginal bacteria and behavioural profiles. Generally, participants who reported limited neovaginal exposures (no penetrative sex, no douching, no dilating, and no application of substances into the neovagina) had the most low-inflammation TC1 and TC4 bacteria, and the least high-inflammation TC2 and TC3 bacteria. Participants who reported diverse exposures, including oral and topical hygienic products, had an opposite profile. However, this does not imply that these exposures are causative. TF may be engaging in diverse activities because of neovaginal discomfort from their microbiota, as opposed to their behaviours causing microbial changes. A potentially interesting observation is that TF who douched and dilated had very similar bacterial profiles to TF who reported only dilating. Vaginal douching is associated with BV and other negative gynecologic and obstetric health outcomes in rCF^79–81^. Many surgical centers recommend douching^82^ to TF after surgery to remove sebum and dilation lubricant (dilation is necessary to maintain neovaginal diameter and depth). However, other centers recommend against it^83^ based on data from rCF. Douching and lubricants can cause osmotic stress in the rCF vagina, which lacks a lipid-rich stratum corneum, and can be detrimental to both the microbiota and the epithelium^84–87^. However, neovaginal tissue has been shown to retain the expression of barrier proteins associated with limiting water loss from skin^42^. Although additional studies are needed, the neovaginal epithelium may not be as sensitive to osmotic stress as the rCF vagina. Additional longitudinal studies powered to assess neovaginal inflammation and microbiota associated with specific douching practices (solution used, frequency, etc.) are warranted. One potentially important contributor to the neovaginal microbiota that was not explored in the current study is sex hormones^88^. Many transfeminine people choose to use exogenous hormones, including estradiol and/or progesterone. Hormone levels were not quantified due to the contactless design of this study; however, it is standard of care to perform orchiectomy at the time of vaginoplasty and all participants reported exogenous estrogen at the time of sample collection. Loss of estrogen after menopause is associated with substantial changes in the vaginal and gut microbiota of females, and androgen deprivation therapy for prostate cancer causes significant changes in the gut microbiota^89–93^. The effect of exogenous estrogen on the neovaginal microbiota and immune milieu should be explored in future *in vivo* and *in vitro* studies.

Conclusions on causality or directionality of the relationships between bacteria and cytokines, symptoms, or behaviours cannot be drawn from this study due to the short observation period (3 weeks), the lack of standardized behaviours and exposures, and the relatively small sample size (n=47). Longer or targeted follow-up with controlled exposures or interventions are needed. In addition, many bacteria found in the neovagina have not been characterized, and taxonomy resolution using sequences of 16S rRNA gene amplicons of the V3-V4 region is limited. Future studies that employ metagenomic sequencing or that characterize neovaginal bacterial isolates would provide new information. A final and important limitation of this study is that it enrolled only participants who had penile inversion vaginoplasty. Other tissues, such as peritoneum or sigmoid colon, may be used to provide neovaginal depth, and based on this study, these tissues would be expected to result in different microbiota.

Despite these limitations, this study can inform current clinical practices. Vaginal pH > 4.5 is considered a clinical indicator of BV in rCF^94^, but neovaginal pH was in the normal range for skin (5.5) and pH correlated positively with the abundance of potentially beneficial TC4 bacteria. Metronidazole is frequently prescribed to TF experiencing BV-like symptoms. This antibiotic is effective against the gram-negative anaerobes that cause BV and – importantly – spares lactobacilli. However, based on the present data, *Lactobacillus* dominance is rarely naturally attainable in the neovagina, and neovaginal cytokines were associated with gram-positive anaerobes, against which metronidazole is expected to have limited efficacy (aside from *Fusobacterium*)^95,96^. While our data calls into question the utility of metronidazole, the overlapping predicted antibiotic sensitivities of low and high inflammation bacteria make selection of an alternative antibiotic difficult. For example, clindamycin and penicillin have activity against gram-positive anaerobes and would be expected to eliminate inflammation-associated genera (*Schaalia*, *Streptococcus*, *Parvimonas*, and *Anaerococcus*), but would also be expected to eliminate potentially beneficial *Ezakiella*^97,98^. *In vitro* studies using neovaginal bacterial isolates may help identify an antibiotic that selectively targets inflammation-associated genera. In addition, such work could lead to the discovery of novel probiotic bacterial strains capable of outcompeting these genera and promoting a low-inflammation environment.

This study represents an initial comprehensive description of the neovaginal microbiota and immune environment. Neovaginal microbiota structure and composition do not resemble the rCF vagina, and it is unlikely that clinical guidelines (both diagnostics and treatments) developed based on research in rCF will be effective for TF. While there is more overlap in microbiota composition between the neovagina and the sub-preputial space of the penis, bacteria associated with inflammation or immunomodulation differ between the two sites. Additional research is needed to develop evidence-based clinical guidelines for transfeminine gynecologic care.

## METHODS

### Study Participants

Transfeminine participants (TF) were a part of the TransBiota study, which characterized the genital microenvironments of trans and gender diverse individuals who have received gender-affirming medical care^60^. Eligible participants were adult (≥18 years old) Canadians that identified as transfeminine and had undergone vaginoplasty ≥ 1 year prior. Participants were recruited online via social media, community groups, healthcare provider referrals, and recontact of consenting TransPULSE Canada participants^99^. Consent was obtained online through REDCap, and consenting TF participants were mailed a study kit with an instruction booklet and materials to self-collect neovaginal samples weekly for three consecutive weeks. Participants were also provided a link to an online questionnaire to be completed at the time of sample collection, capturing demographics, symptoms, and behaviours/exposures that may be relevant to the genital microenvironment^60^.

A cross-sectional cohort of cisgender participants was recruited from London, Ontario. Adult (≥18 years old), uncircumcised, cisgender male (CM, n=56) participants who did not have clinically apparent symptoms of an STI were recruited from the urology clinic at St. Joseph’s Hospital in London, Ontario, and online through social media. Cisgender female (rCF, n=35) participants were recruited online and in person at Western University. Eligible rCF participants were 18-40 years old and provided their sample when not menstruating. Consenting cisgender participants recruited through social media were provided the option to attend Western University for sample collection, or to provide their mailing address for test kits to be sent by mail.

All participants provided written informed consent and institutional research ethics board approval was obtained from Western University (Protocol 115503).

### Sample Collection

At each weekly timepoint, TF participants collected one swab for bacteria characterization, one for immune analyte characterization, and one to evaluate pH. Cisgender participants provided one swab sample for bacteria characterization only. Swabs for bacterial analysis were collected into 1ml of Qiagen C2.1 solution, a validated nucleic acid preservative that affords stabilization of both DNA and RNA molecules at room temperature for up to one month. Swabs for immune analyte quantification were collected into 500μl of phosphate buffered saline (PBS) with protease inhibitor (Complete Mini Protease Inhibitor Cocktail, Sigma-Aldrich), antimicrobial (Primocin, Invivogen), and 10% bovine serum albumin (Sigma-Aldrich).

Participants were instructed to self-collect swabs as follows: (1) TF neovaginal and rCF vaginal: A swab was inserted 5cm into the neovaginal/vaginal canal while avoiding contact with the outer genitalia. The swab was rotated 5 times rubbing the inner neovaginal/vaginal canal wall, removed, and inserted into collection media and swirled for 30 seconds. The swab was then discarded, and the sample tube was sealed shut and shaken for 30 seconds. For TF, this process was repeated for cytokine media, and a third swab was collected, rolled on a pH strip (provided), and the colour of the pH strip was recorded on a provided reference sheet. (2) CM penile: The foreskin was retracted with the non-dominant hand and with the dominant hand a swab was rubbed along the junction between the glans penis and the shaft (the coronal sulcus), back and forth 3 times. The swab was inserted into the collection media and swirled for 30 seconds. The swab was then discarded, and the sample tube was sealed shut and shaken for 30 seconds.

Participants participating by mail (including all TF) returned swab eluants (and pH reference strips, for TF participants) using provided postage-paid envelopes. Swab eluants were aliquoted and stored at -80°C prior to analysis.

### Immune Analyte Quantification

Neovaginal swabs were assessed for concentrations of 19 immune analytes using the Milliplex Cytokine Panel A (GM-CSF, IFN-α2, IFNγ, IL-1α, IL-1β, IL-6, IL-8, IL-10, IL-13, IL-17A, IL-17E/IL-25, IL-17F, IL-22, IP-10, MIG, MIP-1α, MIP-1β, RANTES, IFNγ) on a Luminex MAGPIX system, in accordance with the manufacturer’s instructions (HCYTA-60K-20 Human Cyto Panel A, Millipore Sigma). Plate washing was standardized using an automated plate washer (BioTek 405 TS washer). The lower limit of quantification (LLOQ) was assigned for each analyte based standard curves and using Belysa Immunoassay Curve Fitting Software^100^, which defines the LLOQ as the lowest concentration with CV ≤20% and calculated recovery rate 80-120%.

For three analytes (IL-1β, MIP-1β and IP-10), participant samples measuring at the bottom of the standard curve (i.e., above the Belysa LLOQ but in the lower concentration range) returned poor CV values (>20%). For these analytes, the Belysa LLOQ (based on only the standard curve) was re-designated as the lower limit of detection (LLOD), and a new LLOQ was set as the concentration at which all samples returned a CV >20% (allowing for one re-run of samples with a CV >20%). For analytes with both an LLOD and an LLOQ, samples falling between the LLOD and LLOQ were deemed detectable but not quantifiable, and assigned a concentration corresponding to the midpoint between the LLOD and LLOQ for analytic purposes. Samples with a calculated concentration below the LLOD were assigned a concentration of 0 pg/ml. For all other analytes (IFNγ, IL-1α, IL-6, IL-8, IL-10, IL-22, MIG, MIP-1α, RANTES, IFNγ), the LLOQ corresponded to the lower limit of detection and samples with a calculated concentration below the LLOQ were assigned a concentration of 0 pg/ml.

### Bacterial characterization

Genomic DNA was isolated from swab eluant using the MagAttract PowerMicrobiome DNA/RNA kit (Qiagen) on a liquid handling Microlab STAR (Hamilton). After thawing on ice, 200μl of sample was used as input for a two-step PCR, with dual index barcoding, to amplify the V3-V4 regions of the 16S rRNA gene. Amplicons were pooled at equimolar concentrations, purified, and sequenced on either an Illumina HiSeq 2500 (TF, rCF) or an Illumina NextSeq 1000 (CM) to generate 300-bp paired end reads^101^. Sequences were demultiplexed via a dual barcode strategy, using a mapping file linking barcode to sample ID and split_libraries.py, a QIIME-dependent script^102^. Resulting forward and reverse fastq files were then split by sample, using the QIIME-dependent script split_sequence_file_on_sample_ids.py, and primer sequences were removed via TagCleaner v.0.16^103^. Each amplicon sequencing variant (ASV; 1,684 unique 16S rRNA were obtained from neovaginal swabs) generated by dada2 was assigned taxonomy using the RDP Naïve Bayesian Classifier trained with the SILVA 16S rRNA gene database v138.2^104,105^. Species-level assignments were performed using SpeciateIT v2.0.0^106^. ASVs with matching taxonomic assignments were summed. The core microbiota was determined using the microbiome package in R and was defined as genera present in at least 50% of TF samples (133 samples from 47 individuals) with a relative abundance of at least 1%.

### Statistical Analyses

Shannon diversity index was calculated for TF, rCF and CM groups using the vegan package in R and compared using the Wilcoxon test. The median relative abundances of the top 30 most abundant genera in TF samples (based on median relative abundance) were compared to the rCF and CM groups using the Wilcoxon test. To compare participant demographics across the groups, Fisher’s exact test was used for categorical variables, while continuous variables were compared using the Kruskal-Wallis test.

To identify bacterial taxa that co-occur in vivo in TF samples, Spearman correlations of relative abundance were calculated using the rstatix package in R for the 30 most abundant taxa. Taxa Clusters (TCs) of highly cooccurring bacteria were identified from the Spearman correlation values using the K-means method implemented using the factoextra package with *k* = 4 - as determined using the elbow method - and requiring a minimum of 3 taxa per cluster. Spearman’s correlations were also calculated between taxa versus cytokine concentrations to visualize associations between inflammation and bacterial taxa.

For each sample, we summed the relative abundances of the taxa within each TC to create a TC-weighted dataset. To analyze the association between each TC abundance and age, time since vaginoplasty, pH, behavioural profiles, pre-vaginoplasty circumcision status, and self-reported symptoms, we implemented zero-inflated beta regression models with random effects using the gamlss package to account for repeated measurements from participants. We performed a 5-fold cross-validation to assess the predictive performance and stability of our zero-inflated beta regression models. For categorical variables such as behavioural profiles, pre-vaginoplasty circumcision status, and self-reported symptoms, the categories limited exposures, uncircumcised and no-reported symptom were used as references for comparisons.

The Spearman’s correlation matrix was transformed to a distance matrix using the equation: (1-*cormat*)/2, then hierarchical clustering was performed by hclust and cutreeDynamic into clusters of samples. Immune analyte concentrations were stratified by cluster of samples, and the Kruskal-Wallis test was applied to test for differences in distribution, followed by Dunn’s test with BH correction to test for pairwise differences between groups. The probabilities of transitioning between cluster of samples were calculated and displayed in a heatmap.

Using CLR-transformed 16S rRNA data of all TF study samples (n=133) at genus level and Aitchison distances, a PCA and beta dispersion analysis were performed in order to search for associations between microbiota composition and behavioral profiles as well as self-reported symptoms. Differentially abundant taxa were identified using MaAsLin2 accounting for multiple sampling with default parameters using the relative abundances as response and CS, BC and self-reported symptoms as covariates. All statistical analyses were performed in R Version 4.3.2, and the false discovery rate (fdr) method was used to adjust p-values.

## DATA AVAILABILITY

The 16S rRNA gene amplicon sequence datasets generated during this study are available at NCBI/SRA under accession number PRJNA1230205. All source codes used to analyze the data and generate the figures presented are available in GitHub at github.com/Ravel-lab/TransBiota/ and at github.com/Prodger-Lab/TransBiota/

## ACKNOWLEDGEMENTS

Research reported in this publication was supported by the National Institute of Allergy and Infectious Diseases (NIAID) of the National Institutes of Health (NIH) under award R21AI157912. The Canadian Institutes for Health Research (CIHR PJT 180322), and the CIHR National Women’s Health Research Initiative (NWI 191322). JLP is supported by the Canada Research Chairs Program (CRC-2020-00175). AS is supported by a Canada Graduate Scholarship from CIHR. This research was undertaken, in part, thanks to funding from the Canada Foundation for Innovation (CFI 42343). The authors would like to thank Jason Hallarn and Greta Bauer for their contributions establishing TransBiota. All sequencing was performed by Maryland Genomics (MDG) at the Institute for Genome Sciences, University of Maryland School of Medicine (UMSOM). We acknowledge assistance from Mike Humphrys at MDG with all aspects of sample collection to sequencing.

The funders played no role in study design, data collection, analysis and interpretation of data, or the writing of this manuscript.

## AUTHOR CONTRIBUTIONS

Conceptualization: J.R., J.L.P., Y.K, E.P. Methodology: H.W., B.M., D.Z. Formal analysis: J.R-V., V.T., B.M., H.W., P.G.. Interpretation: J.R-V., H.W., B.M., J.R. and J.L.P. Writing: J.R-V, H.W, J.L.P, Editing: J.R-V., H.W., B.M., P.G., D.Z., A.S., R.P., P.H., V.T., Y.K, E.P., J.L.P. and J.R. Funding acquisition: J.R. and J.L.P.

