## Supplemental Material for "The Neovaginal Microbiota, Symptoms, and Local Immune Correlates in Transfeminine Individuals with Penile Inversion Vaginoplasty"

**Supplemental Table 1. Participant demographics**

| | TF (n=47) | rCF (n=35) | p-value** | CM (n=56) | p-value** |
|---|---|---|---|---|---|
| **Age, in years** (mean, range) | 41.2 (23-69) | 25.1 (20-34) | <0.001 | 30.9 (18-87) | <0.001 |
| **Years since vaginoplasty** (mean, range) | 4.3 (1 - 19) | n/a | - | n/a | - |
| **Circumcised*** (n, %) | 24 (51.1) | n/a | - | 0 (0.0) | |
| **Ethnicity** (n, %) | | | | | |
| White | 42 (89.4) | 19 (54.3) | <0.001 | 28 (50.0) | <0.001 |
| East Asian | 1 (2.1) | 4 (11.4) | | 13 (23.2) | |
| South Asian | 0 | 5 (14.3) | | 5 (9.0) | |
| Middle Eastern | 0 | 3 (8.6) | | 0 | |
| Mixed Ethnicity | 2 (4.3) | 4 (11.4) | | 4 (7.1) | |
| Other | 2 (4.3) | 0 | | 6 (10.7) | |

*For TF, this refers to the pre-vaginoplasty circumcision status.
**Comparing to transfeminine group, calculated using Fisher's exact test (categorical) or Kruskal-Wallis (continuous).

**Supplemental Table 2: Taxa of bacteria identified in the TF neovagina.**

| Taxa | Preva-lence (%) | Relative Abundance (%) | | | | | |
|---|---|---|---|---|---|---|---|
| | | Median | Mean | Min | Q25 | Q75 | Max |
| Absconditabacteriales (SR1) ($\delta$) | 1.504 | 0.000 | 0.001 | 0.000 | 0.000 | 0.000 | 0.109 |
| *Acinetobacter* | 1.504 | 0.000 | 0.001 | 0.000 | 0.000 | 0.000 | 0.097 |
| Actinobacteria ($\gamma$) | 2.256 | 0.000 | 0.000 | 0.000 | 0.000 | 0.000 | 0.007 |
| *Actinobaculum* | 8.271 | 0.000 | 0.040 | 0.000 | 0.000 | 0.000 | 1.736 |
| *Actinobaculum spp.* | 0.752 | 0.000 | 0.000 | 0.000 | 0.000 | 0.000 | 0.015 |
| *Actinobaculum massiliense* | 7.519 | 0.000 | 0.040 | 0.000 | 0.000 | 0.000 | 1.736 |
| *Actinomyces* | 1.504 | 0.000 | 0.001 | 0.000 | 0.000 | 0.000 | 0.048 |
| *Actinomyces urogenitalis* | 1.504 | 0.000 | 0.001 | 0.000 | 0.000 | 0.000 | 0.048 |
| Actinomycetaceae ($\varepsilon$) | 0.752 | 0.000 | 0.000 | 0.000 | 0.000 | 0.000 | 0.006 |
| *Actinotignum* | 60.902 | 0.037 | 0.680 | 0.000 | 0.000 | 0.439 | 9.452 |
| *Actinotignum sanguinis* | 39.098 | 0.000 | 0.310 | 0.000 | 0.000 | 0.077 | 8.520 |
| *Actinotignum schaalii* | 2.256 | 0.000 | 0.007 | 0.000 | 0.000 | 0.000 | 0.562 |
| *Actinotignum timonense* | 25.564 | 0.000 | 0.168 | 0.000 | 0.000 | 0.023 | 9.268 |
| *Actinotignum urinale* | 9.774 | 0.000 | 0.195 | 0.000 | 0.000 | 0.000 | 6.690 |
| *Aerococcus* | 37.594 | 0.000 | 0.297 | 0.000 | 0.000 | 0.117 | 16.311 |
| *Aerococcus christensenii* | 24.060 | 0.000 | 0.227 | 0.000 | 0.000 | 0.000 | 16.311 |
| *Aerococcus sanguinicola* | 4.511 | 0.000 | 0.003 | 0.000 | 0.000 | 0.000 | 0.132 |
| *Aerococcus urinae* | 16.541 | 0.000 | 0.068 | 0.000 | 0.000 | 0.000 | 2.641 |
| *Agathobacter* | 1.504 | 0.000 | 0.000 | 0.000 | 0.000 | 0.000 | 0.039 |
| *Agathobacter rectalis* | 1.504 | 0.000 | 0.000 | 0.000 | 0.000 | 0.000 | 0.039 |
| *Alistipes* | 1.504 | 0.000 | 0.000 | 0.000 | 0.000 | 0.000 | 0.012 |
| *Allisonella* | 0.752 | 0.000 | 0.000 | 0.000 | 0.000 | 0.000 | 0.018 |
| *Allofustis* | 0.752 | 0.000 | 0.000 | 0.000 | 0.000 | 0.000 | 0.007 |
| *Alloiococcus* | 0.752 | 0.000 | 0.000 | 0.000 | 0.000 | 0.000 | 0.035 |
| *Alloprevotella* | 3.759 | 0.000 | 0.004 | 0.000 | 0.000 | 0.000 | 0.308 |
| *Alloscardovia* | 21.053 | 0.000 | 0.136 | 0.000 | 0.000 | 0.000 | 4.773 |
| *Alloscardovia spp.* | 0.752 | 0.000 | 0.000 | 0.000 | 0.000 | 0.000 | 0.033 |
| *Alloscardovia omnicolens* | 21.053 | 0.000 | 0.135 | 0.000 | 0.000 | 0.000 | 4.773 |
| *Anaerobutyricum* | 0.752 | 0.000 | 0.000 | 0.000 | 0.000 | 0.000 | 0.025 |

| | | | | | | | |
|---|---|---|---|---|---|---|---|
| *Anaerobutyricum hallii* | 0.752 | 0.000 | 0.000 | 0.000 | 0.000 | 0.000 | 0.025 |
| *Anaerococcus* | 97.744 | 3.040 | 4.626 | 0.000 | 0.832 | 6.274 | 27.417 |
| *Anaerococcus spp.* | 49.624 | 0.000 | 0.534 | 0.000 | 0.000 | 0.251 | 15.649 |
| *Anaerococcus degeneri* | 5.263 | 0.000 | 0.065 | 0.000 | 0.000 | 0.000 | 3.617 |
| *Anaerococcus hydrogenalis* | 18.797 | 0.000 | 0.067 | 0.000 | 0.000 | 0.000 | 2.848 |
| *Anaerococcus ihuae* | 2.256 | 0.000 | 0.003 | 0.000 | 0.000 | 0.000 | 0.194 |
| *Anaerococcus jeddahensis* | 0.752 | 0.000 | 0.001 | 0.000 | 0.000 | 0.000 | 0.162 |
| *Anaerococcus lactolyticus* | 61.654 | 0.126 | 0.975 | 0.000 | 0.000 | 0.815 | 10.116 |
| *Anaerococcus marasmi* | 21.053 | 0.000 | 0.086 | 0.000 | 0.000 | 0.000 | 5.085 |
| *Anaerococcus murdochii* | 75.940 | 0.491 | 1.298 | 0.000 | 0.000 | 1.723 | 10.802 |
| *Anaerococcus octavius* | 2.256 | 0.000 | 0.001 | 0.000 | 0.000 | 0.000 | 0.083 |
| *Anaerococcus prevotii* | 21.805 | 0.000 | 0.541 | 0.000 | 0.000 | 0.000 | 27.417 |
| *Anaerococcus provencensis* | 4.511 | 0.000 | 0.003 | 0.000 | 0.000 | 0.000 | 0.138 |
| *Anaerococcus rubeinfantis* | 2.256 | 0.000 | 0.007 | 0.000 | 0.000 | 0.000 | 0.803 |
| *Anaerococcus senegalensis* | 6.767 | 0.000 | 0.007 | 0.000 | 0.000 | 0.000 | 0.274 |
| *Anaerococcus vaginalis* | 79.699 | 0.273 | 1.003 | 0.000 | 0.000 | 1.290 | 8.560 |
| *Anaerococcus vaginimassiliensis* | 9.023 | 0.000 | 0.035 | 0.000 | 0.000 | 0.000 | 1.747 |
| *Anaeroglobus* | 40.602 | 0.000 | 0.723 | 0.000 | 0.000 | 0.529 | 12.614 |
| *Anaeroglobus spp.* | 37.594 | 0.000 | 0.501 | 0.000 | 0.000 | 0.153 | 7.581 |
| *Anaeroglobus geminatus* | 5.263 | 0.000 | 0.222 | 0.000 | 0.000 | 0.000 | 12.469 |
| *Anaerostipes* | 0.752 | 0.000 | 0.000 | 0.000 | 0.000 | 0.000 | 0.010 |
| *Anaerostipes hadrus* | 0.752 | 0.000 | 0.000 | 0.000 | 0.000 | 0.000 | 0.010 |
| Anaerovoracaceae (ε) | 5.263 | 0.000 | 0.013 | 0.000 | 0.000 | 0.000 | 0.709 |
| *Aquabacterium* | 0.752 | 0.000 | 0.000 | 0.000 | 0.000 | 0.000 | 0.006 |
| *Arcanobacterium* | 56.391 | 0.030 | 0.243 | 0.000 | 0.000 | 0.367 | 1.618 |
| *Arcanobacterium spp.* | 0.752 | 0.000 | 0.000 | 0.000 | 0.000 | 0.000 | 0.045 |
| *Arcanobacterium urinimassiliense* | 56.391 | 0.030 | 0.240 | 0.000 | 0.000 | 0.349 | 1.618 |

| | | | | | | | |
|---|---|---|---|---|---|---|---|
| *Arcanobacterium pinnipediorum* | 0.752 | 0.000 | 0.003 | 0.000 | 0.000 | 0.000 | 0.455 |
| *Atopobium* | 68.421 | 0.208 | 0.608 | 0.000 | 0.000 | 0.823 | 3.747 |
| *Atopobium spp.* | 36.842 | 0.000 | 0.078 | 0.000 | 0.000 | 0.064 | 1.218 |
| *Atopobium deltae* | 60.150 | 0.100 | 0.524 | 0.000 | 0.000 | 0.706 | 3.747 |
| *Atopobium minutum* | 1.504 | 0.000 | 0.006 | 0.000 | 0.000 | 0.000 | 0.740 |
| Bacilli ($\gamma$) | 1.504 | 0.000 | 0.000 | 0.000 | 0.000 | 0.000 | 0.016 |
| Bacillota ($\beta$) | 0.752 | 0.000 | 0.000 | 0.000 | 0.000 | 0.000 | 0.004 |
| Bacteria ($\alpha$) | 20.301 | 0.000 | 0.012 | 0.000 | 0.000 | 0.000 | 0.355 |
| Bacteroidales ($\delta$) | 18.797 | 0.000 | 0.043 | 0.000 | 0.000 | 0.000 | 1.290 |
| *Bacteroides* | 14.286 | 0.000 | 0.327 | 0.000 | 0.000 | 0.000 | 22.198 |
| *Bacteroides spp.* | 8.271 | 0.000 | 0.007 | 0.000 | 0.000 | 0.000 | 0.266 |
| *Bacteroides fragilis* | 6.015 | 0.000 | 0.320 | 0.000 | 0.000 | 0.000 | 22.198 |
| Bacteroidia ($\gamma$) | 8.271 | 0.000 | 0.010 | 0.000 | 0.000 | 0.000 | 0.346 |
| *Barnesiella* | 1.504 | 0.000 | 0.001 | 0.000 | 0.000 | 0.000 | 0.055 |
| *Barnesiella spp.* | 1.504 | 0.000 | 0.000 | 0.000 | 0.000 | 0.000 | 0.026 |
| *Barnesiella intestinihominis* | 1.504 | 0.000 | 0.000 | 0.000 | 0.000 | 0.000 | 0.029 |
| *Bergeyella* | 1.504 | 0.000 | 0.000 | 0.000 | 0.000 | 0.000 | 0.021 |
| *Berryella* | 17.293 | 0.000 | 0.027 | 0.000 | 0.000 | 0.000 | 0.581 |
| *Bifidobacterium* | 25.564 | 0.000 | 0.160 | 0.000 | 0.000 | 0.010 | 3.129 |
| *Bifidobacterium adolescentis* | 0.752 | 0.000 | 0.000 | 0.000 | 0.000 | 0.000 | 0.018 |
| *Bifidobacterium bifidum* | 0.752 | 0.000 | 0.000 | 0.000 | 0.000 | 0.000 | 0.049 |
| *Bifidobacterium breve* | 7.519 | 0.000 | 0.013 | 0.000 | 0.000 | 0.000 | 1.353 |
| *Bifidobacterium dentium* | 9.774 | 0.000 | 0.076 | 0.000 | 0.000 | 0.000 | 2.953 |
| *Bifidobacterium infantis* | 9.023 | 0.000 | 0.062 | 0.000 | 0.000 | 0.000 | 2.499 |
| *Bifidobacterium scardovii* | 2.256 | 0.000 | 0.008 | 0.000 | 0.000 | 0.000 | 0.696 |
| *Bilophila* | 8.271 | 0.000 | 0.023 | 0.000 | 0.000 | 0.000 | 1.378 |
| *Blautia* | 0.752 | 0.000 | 0.000 | 0.000 | 0.000 | 0.000 | 0.060 |
| *Blautia spp.* | 0.752 | 0.000 | 0.000 | 0.000 | 0.000 | 0.000 | 0.014 |
| *Blautia wexlerae* | 0.752 | 0.000 | 0.000 | 0.000 | 0.000 | 0.000 | 0.046 |
| *Blvii28 wastewater-sludge group* | 3.759 | 0.000 | 0.008 | 0.000 | 0.000 | 0.000 | 0.756 |
| *Brachybacterium* | 0.752 | 0.000 | 0.000 | 0.000 | 0.000 | 0.000 | 0.029 |
| *Bradyrhizobium* | 2.256 | 0.000 | 0.001 | 0.000 | 0.000 | 0.000 | 0.044 |
| *Brevibacterium* | 5.263 | 0.000 | 0.008 | 0.000 | 0.000 | 0.000 | 0.508 |

| | | | | | | | |
|---|---|---|---|---|---|---|---|
| *Brevibacterium mcbrellneri* | 1.504 | 0.000 | 0.003 | 0.000 | 0.000 | 0.000 | 0.227 |
| *Brevibacterium ravenspurgense* | 5.263 | 0.000 | 0.005 | 0.000 | 0.000 | 0.000 | 0.286 |
| *Bulleidia* | 4.511 | 0.000 | 0.012 | 0.000 | 0.000 | 0.000 | 0.777 |
| *Bulleidia spp.* | 0.752 | 0.000 | 0.000 | 0.000 | 0.000 | 0.000 | 0.022 |
| *Bulleidia extructa* | 3.759 | 0.000 | 0.012 | 0.000 | 0.000 | 0.000 | 0.777 |
| Burkholderiales ($\delta$) | 6.015 | 0.000 | 0.010 | 0.000 | 0.000 | 0.000 | 0.814 |
| *Butyrivibrio* | 1.504 | 0.000 | 0.001 | 0.000 | 0.000 | 0.000 | 0.089 |
| *Campylobacter* | 92.481 | 1.129 | 2.032 | 0.000 | 0.000 | 2.501 | 13.526 |
| Campylobacterales ($\delta$) | 0.752 | 0.000 | 0.000 | 0.000 | 0.000 | 0.000 | 0.044 |
| *Candidatus Saccharimonas* | 1.504 | 0.000 | 0.000 | 0.000 | 0.000 | 0.000 | 0.011 |
| *Catenibacterium* | 0.752 | 0.000 | 0.000 | 0.000 | 0.000 | 0.000 | 0.018 |
| *Catonella* | 3.008 | 0.000 | 0.010 | 0.000 | 0.000 | 0.000 | 0.650 |
| *Catonella spp.* | 1.504 | 0.000 | 0.002 | 0.000 | 0.000 | 0.000 | 0.130 |
| *Catonella morbi* | 3.008 | 0.000 | 0.008 | 0.000 | 0.000 | 0.000 | 0.534 |
| *Centipeda* | 4.511 | 0.000 | 0.013 | 0.000 | 0.000 | 0.000 | 0.777 |
| *Christensenellaceae R-7 group* | 22.556 | 0.000 | 0.084 | 0.000 | 0.000 | 0.000 | 1.569 |
| *Cloacibacillus* | 1.504 | 0.000 | 0.004 | 0.000 | 0.000 | 0.000 | 0.353 |
| Clostridia ($\gamma$) | 2.256 | 0.000 | 0.001 | 0.000 | 0.000 | 0.000 | 0.110 |
| *Clostridia UCG-014* | 18.797 | 0.000 | 0.068 | 0.000 | 0.000 | 0.000 | 1.285 |
| *Clostridioides* | 0.752 | 0.000 | 0.000 | 0.000 | 0.000 | 0.000 | 0.007 |
| *Clostridium* | 0.752 | 0.000 | 0.000 | 0.000 | 0.000 | 0.000 | 0.018 |
| *Clostridium innocuum group* | 1.504 | 0.000 | 0.000 | 0.000 | 0.000 | 0.000 | 0.047 |
| *Colidextribacter* | 2.256 | 0.000 | 0.001 | 0.000 | 0.000 | 0.000 | 0.030 |
| *Collinsella* | 1.504 | 0.000 | 0.000 | 0.000 | 0.000 | 0.000 | 0.060 |
| *Collinsella aerofaciens* | 1.504 | 0.000 | 0.000 | 0.000 | 0.000 | 0.000 | 0.060 |
| Coriobacteriales ($\delta$) | 20.301 | 0.000 | 0.015 | 0.000 | 0.000 | 0.000 | 0.631 |
| Corynebacteriaceae ($\varepsilon$) | 0.752 | 0.000 | 0.000 | 0.000 | 0.000 | 0.000 | 0.003 |
| *Corynebacterium* | 57.143 | 0.021 | 0.978 | 0.000 | 0.000 | 0.557 | 16.422 |
| *Corynebacterium spp.* | 34.586 | 0.000 | 0.285 | 0.000 | 0.000 | 0.034 | 10.316 |
| *Corynebacterium amycolatum* | 8.271 | 0.000 | 0.024 | 0.000 | 0.000 | 0.000 | 0.752 |
| *Corynebacterium atypicum* | 0.752 | 0.000 | 0.000 | 0.000 | 0.000 | 0.000 | 0.061 |
| *Corynebacterium aurimucosum* | 12.030 | 0.000 | 0.019 | 0.000 | 0.000 | 0.000 | 0.707 |

| | | | | | | | |
|---|---|---|---|---|---|---|---|
| *Corynebacterium coyleae* | 6.767 | 0.000 | 0.029 | 0.000 | 0.000 | 0.000 | 2.438 |
| *Corynebacterium imitans* | 8.271 | 0.000 | 0.016 | 0.000 | 0.000 | 0.000 | 0.835 |
| *Corynebacterium kefirresidentii* | 22.556 | 0.000 | 0.108 | 0.000 | 0.000 | 0.000 | 2.477 |
| *Corynebacterium mucifaciens* | 2.256 | 0.000 | 0.013 | 0.000 | 0.000 | 0.000 | 1.523 |
| *Corynebacterium pyruviciproducens* | 19.549 | 0.000 | 0.442 | 0.000 | 0.000 | 0.000 | 16.422 |
| *Corynebacterium riegelii* | 3.008 | 0.000 | 0.003 | 0.000 | 0.000 | 0.000 | 0.220 |
| *Corynebacterium striatum* | 10.526 | 0.000 | 0.025 | 0.000 | 0.000 | 0.000 | 1.224 |
| *Corynebacterium tuscaniense* | 3.008 | 0.000 | 0.005 | 0.000 | 0.000 | 0.000 | 0.261 |
| *Corynebacterium urealyticum* | 5.263 | 0.000 | 0.008 | 0.000 | 0.000 | 0.000 | 0.346 |
| *Criibacterium* | 26.316 | 0.000 | 0.053 | 0.000 | 0.000 | 0.022 | 0.653 |
| *Criibacterium spp.* | 1.504 | 0.000 | 0.001 | 0.000 | 0.000 | 0.000 | 0.093 |
| *Criibacterium bergeronii* | 26.316 | 0.000 | 0.052 | 0.000 | 0.000 | 0.022 | 0.653 |
| *Cryptobacterium* | 3.759 | 0.000 | 0.018 | 0.000 | 0.000 | 0.000 | 1.688 |
| *Cryptobacterium curtum* | 3.759 | 0.000 | 0.018 | 0.000 | 0.000 | 0.000 | 1.688 |
| *Cupriavidus* | 1.504 | 0.000 | 0.002 | 0.000 | 0.000 | 0.000 | 0.290 |
| *Cutibacterium* | 9.774 | 0.000 | 0.040 | 0.000 | 0.000 | 0.000 | 3.496 |
| *Cutibacterium acnes* | 6.767 | 0.000 | 0.012 | 0.000 | 0.000 | 0.000 | 0.834 |
| *Cutibacterium avidum* | 3.008 | 0.000 | 0.028 | 0.000 | 0.000 | 0.000 | 3.496 |
| *Defluviitaleaceae UCG-011* | 1.504 | 0.000 | 0.001 | 0.000 | 0.000 | 0.000 | 0.120 |
| *Dermabacter* | 0.752 | 0.000 | 0.001 | 0.000 | 0.000 | 0.000 | 0.169 |
| *Dermabacter hominis* | 0.752 | 0.000 | 0.001 | 0.000 | 0.000 | 0.000 | 0.169 |
| *Dermacoccus* | 0.752 | 0.000 | 0.000 | 0.000 | 0.000 | 0.000 | 0.022 |
| *Desulfovibrio* | 3.759 | 0.000 | 0.008 | 0.000 | 0.000 | 0.000 | 0.430 |
| *Dialister* | 95.489 | 2.358 | 3.221 | 0.000 | 0.000 | 4.647 | 23.268 |
| *Dorea* | 0.752 | 0.000 | 0.000 | 0.000 | 0.000 | 0.000 | 0.013 |
| *Dorea longicatena* | 0.752 | 0.000 | 0.000 | 0.000 | 0.000 | 0.000 | 0.013 |
| *Eggerthella* | 0.752 | 0.000 | 0.000 | 0.000 | 0.000 | 0.000 | 0.033 |
| *Eggerthella lenta* | 0.752 | 0.000 | 0.000 | 0.000 | 0.000 | 0.000 | 0.033 |
| *Eggerthia* | 11.278 | 0.000 | 0.022 | 0.000 | 0.000 | 0.000 | 0.773 |

| | | | | | | | |
|---|---|---|---|---|---|---|---|
| Enterobacterales ($\delta$) | 0.752 | 0.000 | 0.000 | 0.000 | 0.000 | 0.000 | 0.006 |
| *Enterocloster* | 6.767 | 0.000 | 0.006 | 0.000 | 0.000 | 0.000 | 0.372 |
| *Enterocloster citroniae* | 5.263 | 0.000 | 0.006 | 0.000 | 0.000 | 0.000 | 0.372 |
| *Enterocloster clostridioformis* | 1.504 | 0.000 | 0.000 | 0.000 | 0.000 | 0.000 | 0.027 |
| *Enterococcus* | 13.534 | 0.000 | 0.099 | 0.000 | 0.000 | 0.000 | 7.734 |
| *Enterococcus faecalis* | 13.534 | 0.000 | 0.099 | 0.000 | 0.000 | 0.000 | 7.734 |
| Erysipelotrichaceae ($\varepsilon$) | 2.256 | 0.000 | 0.001 | 0.000 | 0.000 | 0.000 | 0.062 |
| *Erysipelotrichaceae UCG-003* | 0.752 | 0.000 | 0.000 | 0.000 | 0.000 | 0.000 | 0.025 |
| *Erysipelotrichaceae UCG-006* | 0.752 | 0.000 | 0.000 | 0.000 | 0.000 | 0.000 | 0.055 |
| *Escherichia* | 6.015 | 0.000 | 0.403 | 0.000 | 0.000 | 0.000 | 24.515 |
| *Escherichia coli* | 6.015 | 0.000 | 0.403 | 0.000 | 0.000 | 0.000 | 24.515 |
| *Eubacterium brachy group* | 8.271 | 0.000 | 0.047 | 0.000 | 0.000 | 0.000 | 3.177 |
| *Eubacterium saphenum group* | 0.752 | 0.000 | 0.001 | 0.000 | 0.000 | 0.000 | 0.109 |
| *Ezakiella* | 84.211 | 1.351 | 3.294 | 0.000 | 0.151 | 4.204 | 27.290 |
| *Ezakiella spp.* | 84.211 | 1.133 | 3.103 | 0.000 | 0.000 | 3.838 | 27.290 |
| *Ezakiella peruensis* | 18.045 | 0.000 | 0.191 | 0.000 | 0.000 | 0.000 | 4.388 |
| *Facklamia* | 47.368 | 0.000 | 0.252 | 0.000 | 0.000 | 0.127 | 4.005 |
| *Facklamia spp.* | 17.293 | 0.000 | 0.105 | 0.000 | 0.000 | 0.000 | 3.931 |
| *Facklamia hominis* | 42.105 | 0.000 | 0.145 | 0.000 | 0.000 | 0.074 | 3.665 |
| *Facklamia languida* | 0.752 | 0.000 | 0.001 | 0.000 | 0.000 | 0.000 | 0.158 |
| *Faecalibacterium* | 1.504 | 0.000 | 0.000 | 0.000 | 0.000 | 0.000 | 0.028 |
| *Faecalibacterium prausnitzii* | 1.504 | 0.000 | 0.000 | 0.000 | 0.000 | 0.000 | 0.028 |
| *Falsiporphyromonas* | 0.752 | 0.000 | 0.000 | 0.000 | 0.000 | 0.000 | 0.027 |
| Family XI ($\gamma$) | 46.617 | 0.000 | 0.924 | 0.000 | 0.000 | 0.735 | 11.104 |
| *Family XIII AD3011 group* | 1.504 | 0.000 | 0.001 | 0.000 | 0.000 | 0.000 | 0.138 |
| *Family XIII UCG-001* | 7.519 | 0.000 | 0.011 | 0.000 | 0.000 | 0.000 | 0.253 |
| *Fannyhessea* | 33.083 | 0.000 | 0.497 | 0.000 | 0.000 | 0.078 | 12.906 |
| *Fannyhessea spp.* | 0.752 | 0.000 | 0.000 | 0.000 | 0.000 | 0.000 | 0.012 |
| *Fannyhessea vaginae* | 33.083 | 0.000 | 0.496 | 0.000 | 0.000 | 0.078 | 12.906 |
| *Fastidiosipila* | 66.917 | 0.190 | 1.120 | 0.000 | 0.000 | 1.118 | 19.387 |
| *Fastidiosipila spp.* | 47.368 | 0.000 | 0.697 | 0.000 | 0.000 | 0.740 | 7.180 |
| *Fastidiosipila sanguinis* | 42.105 | 0.000 | 0.423 | 0.000 | 0.000 | 0.087 | 19.387 |

| | | | | | | | |
|---|---|---|---|---|---|---|---|
| *Fenollaria* | 78.947 | 1.857 | 4.902 | 0.000 | 0.066 | 7.652 | 33.796 |
| *Fenollaria spp.* | 8.271 | 0.000 | 0.005 | 0.000 | 0.000 | 0.000 | 0.269 |
| *Fenollaria massiliensis* | 78.947 | 1.857 | 4.897 | 0.000 | 0.000 | 7.652 | 33.796 |
| *Filifactor* | 1.504 | 0.000 | 0.009 | 0.000 | 0.000 | 0.000 | 0.655 |
| *Filifactor alocis* | 1.504 | 0.000 | 0.009 | 0.000 | 0.000 | 0.000 | 0.655 |
| *Finegoldia* | 87.970 | 0.899 | 3.419 | 0.000 | 0.106 | 3.110 | 41.482 |
| *Finegoldia spp.* | 9.774 | 0.000 | 0.013 | 0.000 | 0.000 | 0.000 | 0.504 |
| *Finegoldia magna* | 87.970 | 0.899 | 3.406 | 0.000 | 0.000 | 3.110 | 41.482 |
| *Fretibacterium* | 4.511 | 0.000 | 0.019 | 0.000 | 0.000 | 0.000 | 1.258 |
| *Fusicatenibacter* | 1.504 | 0.000 | 0.000 | 0.000 | 0.000 | 0.000 | 0.025 |
| *Fusicatenibacter saccharivorans* | 1.504 | 0.000 | 0.000 | 0.000 | 0.000 | 0.000 | 0.025 |
| Fusobacteriaceae (ε) | 2.256 | 0.000 | 0.001 | 0.000 | 0.000 | 0.000 | 0.110 |
| *Fusobacterium* | 76.692 | 1.014 | 4.801 | 0.000 | 0.019 | 7.948 | 32.309 |
| *Fusobacterium spp.* | 9.023 | 0.000 | 0.147 | 0.000 | 0.000 | 0.000 | 5.557 |
| *Fusobacterium animalis* | 64.662 | 0.432 | 3.757 | 0.000 | 0.000 | 5.071 | 32.309 |
| *Fusobacterium nucleatum* | 2.256 | 0.000 | 0.056 | 0.000 | 0.000 | 0.000 | 4.984 |
| *Fusobacterium periodonticum* | 0.752 | 0.000 | 0.000 | 0.000 | 0.000 | 0.000 | 0.048 |
| *Fusobacterium pseudoperiodonticum* | 1.504 | 0.000 | 0.137 | 0.000 | 0.000 | 0.000 | 11.010 |
| *Fusobacterium vincentii* | 12.030 | 0.000 | 0.703 | 0.000 | 0.000 | 0.000 | 19.996 |
| Gammaproteobacteria (γ) | 0.752 | 0.000 | 0.000 | 0.000 | 0.000 | 0.000 | 0.065 |
| *Gardnerella* | 35.338 | 0.000 | 0.823 | 0.000 | 0.000 | 0.121 | 40.490 |
| *Gardnerella vaginalis* | 35.338 | 0.000 | 0.823 | 0.000 | 0.000 | 0.121 | 40.490 |
| *Gemella* | 17.293 | 0.000 | 0.068 | 0.000 | 0.000 | 0.000 | 2.189 |
| *Gemella asaccharolytica* | 13.534 | 0.000 | 0.059 | 0.000 | 0.000 | 0.000 | 2.189 |
| *Gemella morbillorum* | 2.256 | 0.000 | 0.003 | 0.000 | 0.000 | 0.000 | 0.400 |
| *Gemella sanguinis* | 2.256 | 0.000 | 0.006 | 0.000 | 0.000 | 0.000 | 0.673 |
| *Gemmiger* | 0.752 | 0.000 | 0.000 | 0.000 | 0.000 | 0.000 | 0.039 |
| *Georgenia* | 0.752 | 0.000 | 0.000 | 0.000 | 0.000 | 0.000 | 0.017 |
| *Gleimia* | 26.316 | 0.000 | 0.134 | 0.000 | 0.000 | 0.020 | 9.083 |
| *Gleimia europaea* | 26.316 | 0.000 | 0.134 | 0.000 | 0.000 | 0.020 | 9.083 |
| *Granulicatella* | 1.504 | 0.000 | 0.015 | 0.000 | 0.000 | 0.000 | 1.668 |

| | | | | | | | |
|---|---|---|---|---|---|---|---|
| *Granulicatella adiacens* | 0.752 | 0.000 | 0.003 | 0.000 | 0.000 | 0.000 | 0.351 |
| *Granulicatella elegans* | 0.752 | 0.000 | 0.013 | 0.000 | 0.000 | 0.000 | 1.668 |
| *Haemophilus* | 7.519 | 0.000 | 0.045 | 0.000 | 0.000 | 0.000 | 1.979 |
| *Halomonas* | 0.752 | 0.000 | 0.000 | 0.000 | 0.000 | 0.000 | 0.032 |
| *Helcococcus* | 14.286 | 0.000 | 0.042 | 0.000 | 0.000 | 0.000 | 1.358 |
| *Helcococcus spp.* | 9.023 | 0.000 | 0.030 | 0.000 | 0.000 | 0.000 | 1.260 |
| *Helcococcus massiliensis* | 5.263 | 0.000 | 0.010 | 0.000 | 0.000 | 0.000 | 1.038 |
| *Helcococcus sueciensis* | 6.015 | 0.000 | 0.002 | 0.000 | 0.000 | 0.000 | 0.093 |
| *Holdemanella* | 1.504 | 0.000 | 0.000 | 0.000 | 0.000 | 0.000 | 0.035 |
| *Hornefia* | 1.504 | 0.000 | 0.006 | 0.000 | 0.000 | 0.000 | 0.414 |
| *Hornefia spp.* | 1.504 | 0.000 | 0.005 | 0.000 | 0.000 | 0.000 | 0.336 |
| *Hornefia minuta* | 1.504 | 0.000 | 0.001 | 0.000 | 0.000 | 0.000 | 0.078 |
| *Howardella* | 60.902 | 0.025 | 0.083 | 0.000 | 0.000 | 0.086 | 1.454 |
| *Hoylesella* | 96.992 | 7.774 | 9.736 | 0.000 | 3.804 | 13.552 | 39.395 |
| *Hoylesella spp.* | 63.158 | 0.130 | 0.860 | 0.000 | 0.000 | 0.602 | 11.885 |
| *Prevotella buccalis* | 60.150 | 0.142 | 0.669 | 0.000 | 0.000 | 0.545 | 14.531 |
| *Prevotella timonensis* | 88.722 | 5.909 | 8.207 | 0.000 | 0.000 | 11.687 | 38.863 |
| *HT002* | 3.008 | 0.000 | 0.031 | 0.000 | 0.000 | 0.000 | 2.282 |
| *Intestinimonas* | 0.752 | 0.000 | 0.000 | 0.000 | 0.000 | 0.000 | 0.008 |
| *Jonquetella* | 24.812 | 0.000 | 0.650 | 0.000 | 0.000 | 0.000 | 13.399 |
| *Jonquetella spp.* | 5.263 | 0.000 | 0.002 | 0.000 | 0.000 | 0.000 | 0.119 |
| *Jonquetella anthropi* | 24.812 | 0.000 | 0.649 | 0.000 | 0.000 | 0.000 | 13.399 |
| *Kallipyga* | 30.827 | 0.000 | 0.125 | 0.000 | 0.000 | 0.024 | 2.592 |
| *Kallipyga spp.* | 26.316 | 0.000 | 0.123 | 0.000 | 0.000 | 0.018 | 2.592 |
| *Kallipyga gabonensis* | 4.511 | 0.000 | 0.002 | 0.000 | 0.000 | 0.000 | 0.134 |
| *Kocuria* | 0.752 | 0.000 | 0.001 | 0.000 | 0.000 | 0.000 | 0.137 |
| *Lachnoanaerobaculum* | 3.008 | 0.000 | 0.009 | 0.000 | 0.000 | 0.000 | 0.797 |
| *Lachnoanaerobaculum spp.* | 2.256 | 0.000 | 0.008 | 0.000 | 0.000 | 0.000 | 0.797 |
| *Lachnoanaerobaculum orale* | 0.752 | 0.000 | 0.000 | 0.000 | 0.000 | 0.000 | 0.048 |
| *Lachnoclostridium* | 0.752 | 0.000 | 0.000 | 0.000 | 0.000 | 0.000 | 0.025 |
| *Lachnospira* | 0.752 | 0.000 | 0.000 | 0.000 | 0.000 | 0.000 | 0.010 |
| *Lachnospira rogosae* | 0.752 | 0.000 | 0.000 | 0.000 | 0.000 | 0.000 | 0.010 |
| Lachnospiraceae ($\varepsilon$) | 1.504 | 0.000 | 0.000 | 0.000 | 0.000 | 0.000 | 0.021 |
| *Lachnospiraceae FE2018 group* | 9.774 | 0.000 | 0.011 | 0.000 | 0.000 | 0.000 | 0.315 |

| | | | | | | | |
|---|---|---|---|---|---|---|---|
| *Lachnospiraceae ND3007 group* | 0.752 | 0.000 | 0.000 | 0.000 | 0.000 | 0.000 | 0.014 |
| *Lachnospiraceae NK4A136 group* | 0.752 | 0.000 | 0.000 | 0.000 | 0.000 | 0.000 | 0.010 |
| *Lachnospiraceae UCG-010* | 0.752 | 0.000 | 0.000 | 0.000 | 0.000 | 0.000 | 0.008 |
| *Lacticaseibacillus* | 1.504 | 0.000 | 0.001 | 0.000 | 0.000 | 0.000 | 0.112 |
| *Lacticaseibacillus rhamnosus* | 1.504 | 0.000 | 0.001 | 0.000 | 0.000 | 0.000 | 0.112 |
| Lactobacillales ($\delta$) | 2.256 | 0.000 | 0.001 | 0.000 | 0.000 | 0.000 | 0.052 |
| *Lactobacillus* | 60.902 | 0.074 | 2.359 | 0.000 | 0.000 | 0.316 | 72.576 |
| *Lactobacillus spp.* | 1.504 | 0.000 | 0.002 | 0.000 | 0.000 | 0.000 | 0.213 |
| *Lactobacillus crispatus* | 20.301 | 0.000 | 0.172 | 0.000 | 0.000 | 0.000 | 16.264 |
| *Lactobacillus gasseri* | 15.789 | 0.000 | 1.345 | 0.000 | 0.000 | 0.000 | 71.908 |
| *Lactobacillus iners* | 45.865 | 0.000 | 0.658 | 0.000 | 0.000 | 0.132 | 14.805 |
| *Lactobacillus jensenii* | 7.519 | 0.000 | 0.179 | 0.000 | 0.000 | 0.000 | 15.988 |
| *Lactobacillus mulieris* | 1.504 | 0.000 | 0.000 | 0.000 | 0.000 | 0.000 | 0.039 |
| *Lactobacillus paragasseri* | 0.752 | 0.000 | 0.003 | 0.000 | 0.000 | 0.000 | 0.363 |
| *Lancefieldella* | 24.060 | 0.000 | 0.156 | 0.000 | 0.000 | 0.000 | 3.836 |
| *Lancefieldella spp.* | 3.759 | 0.000 | 0.015 | 0.000 | 0.000 | 0.000 | 1.078 |
| *Lancefieldella parvula* | 6.015 | 0.000 | 0.022 | 0.000 | 0.000 | 0.000 | 1.274 |
| *Lancefieldella rimae* | 21.053 | 0.000 | 0.118 | 0.000 | 0.000 | 0.000 | 3.325 |
| *Lawsonella* | 91.729 | 0.336 | 1.266 | 0.000 | 0.086 | 0.826 | 32.772 |
| *Lawsonella spp.* | 2.256 | 0.000 | 0.002 | 0.000 | 0.000 | 0.000 | 0.107 |
| *Lawsonella clevelandensis* | 91.729 | 0.336 | 1.265 | 0.000 | 0.000 | 0.826 | 32.772 |
| Leptotrichiaceae ($\varepsilon$) | 0.752 | 0.000 | 0.000 | 0.000 | 0.000 | 0.000 | 0.056 |
| *Ligilactobacillus* | 0.752 | 0.000 | 0.001 | 0.000 | 0.000 | 0.000 | 0.162 |
| *Ligilactobacillus salivarius* | 0.752 | 0.000 | 0.001 | 0.000 | 0.000 | 0.000 | 0.162 |
| *Limosilactobacillus* | 4.511 | 0.000 | 0.113 | 0.000 | 0.000 | 0.000 | 11.802 |
| *Limosilactobacillus coleohominis* | 3.008 | 0.000 | 0.030 | 0.000 | 0.000 | 0.000 | 2.483 |
| *Limosilactobacillus fermentum* | 1.504 | 0.000 | 0.081 | 0.000 | 0.000 | 0.000 | 10.342 |
| *Limosilactobacillus oris* | 0.752 | 0.000 | 0.000 | 0.000 | 0.000 | 0.000 | 0.054 |
| *Limosilactobacillus vaginalis* | 1.504 | 0.000 | 0.001 | 0.000 | 0.000 | 0.000 | 0.056 |

| | | | | | | | |
|---|---|---|---|---|---|---|---|
| *Mediterraneibacter* | 1.504 | 0.000 | 0.000 | 0.000 | 0.000 | 0.000 | 0.021 |
| *Mediterraneibacter lactaris* | 0.752 | 0.000 | 0.000 | 0.000 | 0.000 | 0.000 | 0.018 |
| *Mediterraneibacter torques* | 0.752 | 0.000 | 0.000 | 0.000 | 0.000 | 0.000 | 0.021 |
| *Megasphaera* | 18.045 | 0.000 | 0.058 | 0.000 | 0.000 | 0.000 | 1.882 |
| *Megasphaera spp.* | 16.541 | 0.000 | 0.056 | 0.000 | 0.000 | 0.000 | 1.882 |
| *Megasphaera lornae* | 1.504 | 0.000 | 0.002 | 0.000 | 0.000 | 0.000 | 0.243 |
| *Methylobacterium* | 0.752 | 0.000 | 0.000 | 0.000 | 0.000 | 0.000 | 0.014 |
| *Micrococcus* | 0.752 | 0.000 | 0.001 | 0.000 | 0.000 | 0.000 | 0.161 |
| *Millisia* | 0.752 | 0.000 | 0.000 | 0.000 | 0.000 | 0.000 | 0.020 |
| *Mobiluncus* | 78.947 | 0.433 | 1.220 | 0.000 | 0.069 | 1.506 | 14.453 |
| *Mobiluncus spp.* | 2.256 | 0.000 | 0.001 | 0.000 | 0.000 | 0.000 | 0.082 |
| *Mobiluncus curtisii* | 77.444 | 0.398 | 1.193 | 0.000 | 0.000 | 1.439 | 14.453 |
| *Mobiluncus mulieris* | 1.504 | 0.000 | 0.026 | 0.000 | 0.000 | 0.000 | 2.075 |
| *Mogibacterium* | 36.090 | 0.000 | 0.126 | 0.000 | 0.000 | 0.052 | 4.934 |
| *Mogibacterium spp.* | 27.820 | 0.000 | 0.065 | 0.000 | 0.000 | 0.013 | 1.891 |
| *Mogibacterium diversum* | 10.526 | 0.000 | 0.061 | 0.000 | 0.000 | 0.000 | 4.710 |
| *Moryella* | 46.617 | 0.000 | 0.556 | 0.000 | 0.000 | 0.637 | 5.243 |
| *Murdochiella* | 75.940 | 0.246 | 1.481 | 0.000 | 0.027 | 1.707 | 14.080 |
| *Murdochiella spp.* | 68.421 | 0.183 | 1.385 | 0.000 | 0.000 | 1.264 | 14.080 |
| *Murdochiella massiliensis* | 2.256 | 0.000 | 0.001 | 0.000 | 0.000 | 0.000 | 0.069 |
| *Murdochiella vaginalis* | 18.045 | 0.000 | 0.095 | 0.000 | 0.000 | 0.000 | 2.710 |
| Muribaculaceae ($\varepsilon$) | 39.850 | 0.000 | 1.063 | 0.000 | 0.000 | 0.208 | 17.857 |
| Mycobacteriales ($\delta$) | 0.752 | 0.000 | 0.000 | 0.000 | 0.000 | 0.000 | 0.011 |
| *Mycoplasma* | 5.263 | 0.000 | 0.023 | 0.000 | 0.000 | 0.000 | 1.007 |
| *Mycoplasmoides* | 0.752 | 0.000 | 0.000 | 0.000 | 0.000 | 0.000 | 0.019 |
| *Mycoplasmopsis* | 2.256 | 0.000 | 0.009 | 0.000 | 0.000 | 0.000 | 0.687 |
| *Nakamurella* | 0.752 | 0.000 | 0.000 | 0.000 | 0.000 | 0.000 | 0.065 |
| *Ndongobacter* | 18.045 | 0.000 | 0.014 | 0.000 | 0.000 | 0.000 | 0.333 |
| *Ndongobacter massiliensis* | 18.045 | 0.000 | 0.014 | 0.000 | 0.000 | 0.000 | 0.333 |
| *Negativicoccus* | 51.128 | 0.010 | 0.427 | 0.000 | 0.000 | 0.326 | 3.666 |
| *Negativicoccus spp.* | 48.872 | 0.000 | 0.415 | 0.000 | 0.000 | 0.243 | 3.666 |
| *Negativicoccus massiliensis* | 6.767 | 0.000 | 0.012 | 0.000 | 0.000 | 0.000 | 0.754 |
| *Neisseria* | 0.752 | 0.000 | 0.001 | 0.000 | 0.000 | 0.000 | 0.084 |
| *Neisseria gonorrhoeae* | 0.752 | 0.000 | 0.001 | 0.000 | 0.000 | 0.000 | 0.084 |

| | | | | | | | |
|---|---|---|---|---|---|---|---|
| Neisseriaceae ($\varepsilon$) | 0.752 | 0.000 | 0.000 | 0.000 | 0.000 | 0.000 | 0.008 |
| *Nosocomiicoccus* | 6.015 | 0.000 | 0.009 | 0.000 | 0.000 | 0.000 | 0.735 |
| *Nosocomiicoccus spp.* | 5.263 | 0.000 | 0.008 | 0.000 | 0.000 | 0.000 | 0.735 |
| *Nosocomiicoccus ampullae A* | 1.504 | 0.000 | 0.001 | 0.000 | 0.000 | 0.000 | 0.071 |
| *Oligella* | 0.752 | 0.000 | 0.000 | 0.000 | 0.000 | 0.000 | 0.008 |
| *Oligella urethralis* | 0.752 | 0.000 | 0.000 | 0.000 | 0.000 | 0.000 | 0.008 |
| *Olsenella* | 8.271 | 0.000 | 0.009 | 0.000 | 0.000 | 0.000 | 0.357 |
| *Olsenella spp.* | 2.256 | 0.000 | 0.000 | 0.000 | 0.000 | 0.000 | 0.022 |
| *Olsenella uli* | 6.015 | 0.000 | 0.009 | 0.000 | 0.000 | 0.000 | 0.357 |
| *Oribacterium* | 5.263 | 0.000 | 0.008 | 0.000 | 0.000 | 0.000 | 0.329 |
| *Oscillibacter* | 2.256 | 0.000 | 0.000 | 0.000 | 0.000 | 0.000 | 0.018 |
| Oscillospiraceae ($\varepsilon$) | 16.541 | 0.000 | 0.026 | 0.000 | 0.000 | 0.000 | 0.511 |
| *Paracoccus* | 0.752 | 0.000 | 0.000 | 0.000 | 0.000 | 0.000 | 0.017 |
| *Parasutterella* | 1.504 | 0.000 | 0.003 | 0.000 | 0.000 | 0.000 | 0.269 |
| *Parvimonas* | 70.677 | 0.442 | 1.890 | 0.000 | 0.000 | 2.711 | 20.211 |
| *Parvimonas spp.* | 20.301 | 0.000 | 0.323 | 0.000 | 0.000 | 0.000 | 17.157 |
| *Parvimonas micra* | 22.556 | 0.000 | 0.308 | 0.000 | 0.000 | 0.000 | 7.160 |
| *Parvimonas parva* | 45.865 | 0.000 | 1.260 | 0.000 | 0.000 | 0.973 | 20.211 |
| *Pasteurella* | 0.752 | 0.000 | 0.001 | 0.000 | 0.000 | 0.000 | 0.121 |
| *Pauljensenia* | 0.752 | 0.000 | 0.001 | 0.000 | 0.000 | 0.000 | 0.189 |
| *Peptococcus* | 53.383 | 0.033 | 0.480 | 0.000 | 0.000 | 0.444 | 6.216 |
| *Peptoniphilus* | 98.496 | 10.684 | 11.125 | 0.000 | 6.676 | 14.257 | 38.879 |
| *Peptoniphilus spp.* | 88.722 | 2.473 | 3.354 | 0.000 | 0.000 | 4.709 | 19.993 |
| *Peptoniphilus catoniae* | 3.008 | 0.000 | 0.001 | 0.000 | 0.000 | 0.000 | 0.093 |
| *Peptoniphilus coxii* | 62.406 | 0.279 | 0.998 | 0.000 | 0.000 | 1.487 | 9.860 |
| *Peptoniphilus gorbachii* | 19.549 | 0.000 | 0.103 | 0.000 | 0.000 | 0.000 | 2.049 |
| *Peptoniphilus harei* | 96.241 | 1.727 | 2.690 | 0.000 | 0.000 | 4.098 | 17.431 |
| *Peptoniphilus lacrimalis* | 86.466 | 1.809 | 2.230 | 0.000 | 0.000 | 3.561 | 10.575 |
| *Peptoniphilus nemausensis* | 3.759 | 0.000 | 0.005 | 0.000 | 0.000 | 0.000 | 0.219 |
| *Peptoniphilus obesi* | 46.617 | 0.000 | 0.078 | 0.000 | 0.000 | 0.076 | 1.025 |
| *Peptoniphilus pacaensis* | 31.579 | 0.000 | 0.396 | 0.000 | 0.000 | 0.304 | 4.998 |
| *Peptoniphilus phoceensis* | 2.256 | 0.000 | 0.044 | 0.000 | 0.000 | 0.000 | 3.617 |
| *Peptoniphilus rachelemmaiella* | 55.639 | 0.136 | 1.225 | 0.000 | 0.000 | 1.057 | 14.713 |

| | | | | | | | |
|---|---|---|---|---|---|---|---|
| Peptostreptococcales-Tissierellales ($\delta$) | 22.556 | 0.000 | 0.126 | 0.000 | 0.000 | 0.000 | 3.300 |
| *Peptostreptococcus* | 48.120 | 0.000 | 0.984 | 0.000 | 0.000 | 1.424 | 8.683 |
| *Peptostreptococcus spp.* | 16.541 | 0.000 | 0.339 | 0.000 | 0.000 | 0.000 | 8.683 |
| *Peptostreptococcus anaerobius* | 35.338 | 0.000 | 0.646 | 0.000 | 0.000 | 0.158 | 8.284 |
| *Peredibacter* | 0.752 | 0.000 | 0.000 | 0.000 | 0.000 | 0.000 | 0.011 |
| Porphyromonadaceae ($\varepsilon$) | 6.767 | 0.000 | 0.004 | 0.000 | 0.000 | 0.000 | 0.238 |
| *Porphyromonas* | 92.481 | 6.757 | 8.277 | 0.000 | 1.957 | 12.808 | 42.152 |
| *Porphyromonas spp.* | 87.218 | 3.006 | 4.280 | 0.000 | 0.000 | 6.374 | 23.167 |
| *Porphyromonas asaccharolytica* | 70.677 | 0.420 | 1.685 | 0.000 | 0.000 | 2.044 | 11.569 |
| *Porphyromonas gingivalis* | 2.256 | 0.000 | 0.478 | 0.000 | 0.000 | 0.000 | 34.095 |
| *Porphyromonas uenonis* | 67.669 | 0.622 | 1.833 | 0.000 | 0.000 | 2.395 | 21.667 |
| *Prevotella* | 97.744 | 7.778 | 9.235 | 0.000 | 2.066 | 12.195 | 44.355 |
| *Prevotella spp.* | 86.466 | 0.716 | 2.568 | 0.000 | 0.000 | 3.622 | 25.737 |
| *Prevotella amnii* | 2.256 | 0.000 | 0.079 | 0.000 | 0.000 | 0.000 | 6.885 |
| *Prevotella bivia* | 40.602 | 0.000 | 3.556 | 0.000 | 0.000 | 0.553 | 44.202 |
| *Prevotella brunnea* | 1.504 | 0.000 | 0.021 | 0.000 | 0.000 | 0.000 | 1.938 |
| *Prevotella corporis* | 26.316 | 0.000 | 0.990 | 0.000 | 0.000 | 0.014 | 25.352 |
| *Prevotella disiens* | 50.376 | 0.007 | 1.989 | 0.000 | 0.000 | 1.930 | 19.571 |
| *Prevotella intermedia* | 1.504 | 0.000 | 0.012 | 0.000 | 0.000 | 0.000 | 1.568 |
| *Prevotella jejuni* | 0.752 | 0.000 | 0.000 | 0.000 | 0.000 | 0.000 | 0.040 |
| *Prevotella melaninogenica* | 3.008 | 0.000 | 0.021 | 0.000 | 0.000 | 0.000 | 2.189 |
| Prevotellaceae ($\varepsilon$) | 8.271 | 0.000 | 0.033 | 0.000 | 0.000 | 0.000 | 0.749 |
| *Prevotellaceae UCG-001* | 13.534 | 0.000 | 0.141 | 0.000 | 0.000 | 0.000 | 5.195 |
| Propionibacteriaceae ($\varepsilon$) | 1.504 | 0.000 | 0.013 | 0.000 | 0.000 | 0.000 | 1.648 |
| *Propionibacterium* | 14.286 | 0.000 | 0.193 | 0.000 | 0.000 | 0.000 | 8.042 |
| *Propionimicrobium* | 67.669 | 0.058 | 0.816 | 0.000 | 0.000 | 0.498 | 18.178 |
| *Propionimicrobium spp.* | 4.511 | 0.000 | 0.130 | 0.000 | 0.000 | 0.000 | 9.795 |
| *Propionimicrobium lymphophilum* | 66.165 | 0.052 | 0.686 | 0.000 | 0.000 | 0.423 | 18.178 |
| *Proteus* | 1.504 | 0.000 | 0.006 | 0.000 | 0.000 | 0.000 | 0.842 |
| *Pseudoglutamicibacter* | 2.256 | 0.000 | 0.001 | 0.000 | 0.000 | 0.000 | 0.102 |
| *Pseudoglutamicibacter albus* | 2.256 | 0.000 | 0.001 | 0.000 | 0.000 | 0.000 | 0.102 |

| | | | | | | | |
|---|---|---|---|---|---|---|---|
| Pseudomonadota (*β*) | 3.008 | 0.000 | 0.001 | 0.000 | 0.000 | 0.000 | 0.059 |
| *Pseudomonas* | 0.752 | 0.000 | 0.000 | 0.000 | 0.000 | 0.000 | 0.007 |
| *Pseudoramibacter* | 7.519 | 0.000 | 0.011 | 0.000 | 0.000 | 0.000 | 0.646 |
| *Pyramidobacter* | 1.504 | 0.000 | 0.001 | 0.000 | 0.000 | 0.000 | 0.088 |
| *Pyramidobacter piscolens* | 1.504 | 0.000 | 0.001 | 0.000 | 0.000 | 0.000 | 0.088 |
| *Ralstonia* | 0.752 | 0.000 | 0.000 | 0.000 | 0.000 | 0.000 | 0.006 |
| *Rarimicrobium* | 11.278 | 0.000 | 0.065 | 0.000 | 0.000 | 0.000 | 2.487 |
| RF39 (*δ*) | 6.015 | 0.000 | 0.013 | 0.000 | 0.000 | 0.000 | 0.822 |
| Rickettsiales (*δ*) | 0.752 | 0.000 | 0.000 | 0.000 | 0.000 | 0.000 | 0.006 |
| *Rikenellaceae RC9 gut group* | 27.820 | 0.000 | 0.101 | 0.000 | 0.000 | 0.020 | 1.389 |
| *Roseateles* | 0.752 | 0.000 | 0.000 | 0.000 | 0.000 | 0.000 | 0.016 |
| *Rothia* | 2.256 | 0.000 | 0.002 | 0.000 | 0.000 | 0.000 | 0.266 |
| *Rothia aeria* | 2.256 | 0.000 | 0.002 | 0.000 | 0.000 | 0.000 | 0.266 |
| Ruminococcaceae (*ε*) | 0.752 | 0.000 | 0.001 | 0.000 | 0.000 | 0.000 | 0.113 |
| *Ruminococcus* | 0.752 | 0.000 | 0.000 | 0.000 | 0.000 | 0.000 | 0.023 |
| *Ruminococcus gnavus group* | 0.752 | 0.000 | 0.000 | 0.000 | 0.000 | 0.000 | 0.014 |
| *Ruthenibacterium* | 0.752 | 0.000 | 0.000 | 0.000 | 0.000 | 0.000 | 0.017 |
| *Ruthenibacterium lactatiformans* | 0.752 | 0.000 | 0.000 | 0.000 | 0.000 | 0.000 | 0.017 |
| *S5-A14a* | 57.895 | 0.039 | 0.323 | 0.000 | 0.000 | 0.332 | 3.508 |
| Saccharimonadales (*δ*) | 20.301 | 0.000 | 0.077 | 0.000 | 0.000 | 0.000 | 1.538 |
| *Schaalia* | 80.451 | 0.174 | 0.391 | 0.000 | 0.000 | 0.483 | 2.945 |
| *Segatella* | 15.789 | 0.000 | 0.194 | 0.000 | 0.000 | 0.000 | 4.838 |
| *Segatella spp.* | 6.015 | 0.000 | 0.030 | 0.000 | 0.000 | 0.000 | 2.541 |
| *Segatella copri* | 3.759 | 0.000 | 0.007 | 0.000 | 0.000 | 0.000 | 0.412 |
| *Segatella oris* | 11.278 | 0.000 | 0.158 | 0.000 | 0.000 | 0.000 | 4.838 |
| *Selenomonas* | 6.767 | 0.000 | 0.015 | 0.000 | 0.000 | 0.000 | 0.683 |
| *Selenomonas spp.* | 6.015 | 0.000 | 0.015 | 0.000 | 0.000 | 0.000 | 0.683 |
| *Selenomonas sputigena* | 0.752 | 0.000 | 0.000 | 0.000 | 0.000 | 0.000 | 0.062 |
| *Serratia* | 2.256 | 0.000 | 0.008 | 0.000 | 0.000 | 0.000 | 0.982 |
| *Slackia* | 48.872 | 0.000 | 0.084 | 0.000 | 0.000 | 0.073 | 1.373 |
| *Slackia exigua* | 48.872 | 0.000 | 0.084 | 0.000 | 0.000 | 0.073 | 1.373 |
| *Sneathia* | 27.068 | 0.000 | 1.158 | 0.000 | 0.000 | 0.031 | 22.477 |
| *Sneathia spp.* | 21.053 | 0.000 | 0.807 | 0.000 | 0.000 | 0.000 | 22.477 |
| *Sneathia vaginalis* | 12.782 | 0.000 | 0.352 | 0.000 | 0.000 | 0.000 | 9.608 |
| *Solobacterium* | 25.564 | 0.000 | 0.211 | 0.000 | 0.000 | 0.009 | 6.789 |

| | | | | | | | |
|---|---|---|---|---|---|---|---|
| *Staphylococcus* | 26.316 | 0.000 | 0.367 | 0.000 | 0.000 | 0.015 | 20.278 |
| *Staphylococcus spp.* | 0.752 | 0.000 | 0.001 | 0.000 | 0.000 | 0.000 | 0.104 |
| *Staphylococcus epidermidis* | 17.293 | 0.000 | 0.145 | 0.000 | 0.000 | 0.000 | 5.927 |
| *Staphylococcus haemolyticus* | 5.263 | 0.000 | 0.003 | 0.000 | 0.000 | 0.000 | 0.192 |
| *Staphylococcus hominis* | 3.759 | 0.000 | 0.207 | 0.000 | 0.000 | 0.000 | 17.558 |
| *Staphylococcus lugdunensis* | 3.759 | 0.000 | 0.008 | 0.000 | 0.000 | 0.000 | 0.440 |
| *Staphylococcus warneri* | 4.511 | 0.000 | 0.003 | 0.000 | 0.000 | 0.000 | 0.121 |
| *Stomatobaculum* | 0.752 | 0.000 | 0.000 | 0.000 | 0.000 | 0.000 | 0.024 |
| *Streptococcus* | 81.955 | 0.381 | 3.129 | 0.000 | 0.014 | 2.711 | 66.089 |
| *Streptococcus spp.* | 0.752 | 0.000 | 0.000 | 0.000 | 0.000 | 0.000 | 0.006 |
| *Streptococcus agalactiae* | 8.271 | 0.000 | 0.498 | 0.000 | 0.000 | 0.000 | 61.616 |
| *Streptococcus anginosus* | 70.677 | 0.132 | 2.413 | 0.000 | 0.000 | 2.118 | 37.454 |
| *Streptococcus constellatus* | 10.526 | 0.000 | 0.150 | 0.000 | 0.000 | 0.000 | 9.079 |
| *Streptococcus cristatus* | 1.504 | 0.000 | 0.004 | 0.000 | 0.000 | 0.000 | 0.297 |
| *Streptococcus intermedius* | 0.752 | 0.000 | 0.001 | 0.000 | 0.000 | 0.000 | 0.095 |
| *Streptococcus mitis* | 13.534 | 0.000 | 0.062 | 0.000 | 0.000 | 0.000 | 3.358 |
| *Streptococcus salivarius* | 2.256 | 0.000 | 0.002 | 0.000 | 0.000 | 0.000 | 0.266 |
| *Streptococcus urinalis* | 0.752 | 0.000 | 0.000 | 0.000 | 0.000 | 0.000 | 0.023 |
| *Succiniclasticum* | 9.023 | 0.000 | 0.021 | 0.000 | 0.000 | 0.000 | 1.764 |
| *Sutterella* | 24.812 | 0.000 | 0.033 | 0.000 | 0.000 | 0.000 | 0.400 |
| *Sutterella spp.* | 23.308 | 0.000 | 0.030 | 0.000 | 0.000 | 0.000 | 0.394 |
| *Sutterella wadsworthensis* | 2.256 | 0.000 | 0.003 | 0.000 | 0.000 | 0.000 | 0.382 |
| *Tannerella* | 3.008 | 0.000 | 0.004 | 0.000 | 0.000 | 0.000 | 0.281 |
| *Tannerella spp.* | 1.504 | 0.000 | 0.001 | 0.000 | 0.000 | 0.000 | 0.122 |
| *Tannerella forsythia* | 3.008 | 0.000 | 0.003 | 0.000 | 0.000 | 0.000 | 0.159 |
| *Tessaracoccus* | 0.752 | 0.000 | 0.000 | 0.000 | 0.000 | 0.000 | 0.053 |
| *Thomasclavelia* | 3.008 | 0.000 | 0.014 | 0.000 | 0.000 | 0.000 | 1.502 |
| *Treponema* | 12.782 | 0.000 | 0.058 | 0.000 | 0.000 | 0.000 | 2.347 |
| *Trueperella* | 7.519 | 0.000 | 0.008 | 0.000 | 0.000 | 0.000 | 0.268 |
| *Trueperella spp.* | 1.504 | 0.000 | 0.000 | 0.000 | 0.000 | 0.000 | 0.022 |

| | | | | | | | |
|---|---|---|---|---|---|---|---|
| *Trueperella bernardiae* | 6.015 | 0.000 | 0.008 | 0.000 | 0.000 | 0.000 | 0.268 |
| *Ureaplasma* | 3.759 | 0.000 | 0.003 | 0.000 | 0.000 | 0.000 | 0.338 |
| *Ureaplasma parvum* | 3.759 | 0.000 | 0.003 | 0.000 | 0.000 | 0.000 | 0.338 |
| *Varibaculum* | 90.977 | 1.187 | 3.603 | 0.000 | 0.202 | 5.236 | 30.927 |
| *Varibaculum spp.* | 24.812 | 0.000 | 0.466 | 0.000 | 0.000 | 0.000 | 12.348 |
| *Varibaculum cambriense* | 62.406 | 0.092 | 1.318 | 0.000 | 0.000 | 0.806 | 30.701 |
| *Varibaculum massiliense* | 78.947 | 0.321 | 1.819 | 0.000 | 0.000 | 2.072 | 16.315 |
| *Veillonella* | 23.308 | 0.000 | 0.233 | 0.000 | 0.000 | 0.000 | 13.411 |
| *Veillonella spp.* | 0.752 | 0.000 | 0.003 | 0.000 | 0.000 | 0.000 | 0.419 |
| *Veillonella atypica* | 7.519 | 0.000 | 0.093 | 0.000 | 0.000 | 0.000 | 5.139 |
| *Veillonella dispar* | 6.015 | 0.000 | 0.068 | 0.000 | 0.000 | 0.000 | 5.073 |
| *Veillonella montpellierensis* | 0.752 | 0.000 | 0.004 | 0.000 | 0.000 | 0.000 | 0.573 |
| *Veillonella parvula* | 15.038 | 0.000 | 0.065 | 0.000 | 0.000 | 0.000 | 3.724 |
| Veillonellaceae ($\varepsilon$) | 89.474 | 0.481 | 0.649 | 0.000 | 0.000 | 0.762 | 4.580 |
| *Vibrionimonas* | 0.752 | 0.000 | 0.000 | 0.000 | 0.000 | 0.000 | 0.026 |
| *W5053* | 51.128 | 0.042 | 0.530 | 0.000 | 0.000 | 0.786 | 3.853 |
| Williamwhitmaniaceae ($\varepsilon$) | 0.752 | 0.000 | 0.000 | 0.000 | 0.000 | 0.000 | 0.011 |
| *Winkia* | 23.308 | 0.000 | 0.247 | 0.000 | 0.000 | 0.000 | 13.999 |
| *Winkia spp.* | 12.782 | 0.000 | 0.018 | 0.000 | 0.000 | 0.000 | 0.648 |
| *Winkia neuii* | 12.782 | 0.000 | 0.229 | 0.000 | 0.000 | 0.000 | 13.999 |

Taxonomy levels: Domain ($\boldsymbol{\alpha}$), Phylum ($\boldsymbol{\beta}$), Class ($\boldsymbol{\gamma}$), Order ($\boldsymbol{\delta}$), and Family ($\boldsymbol{\varepsilon}$)
**Q:** Quartile

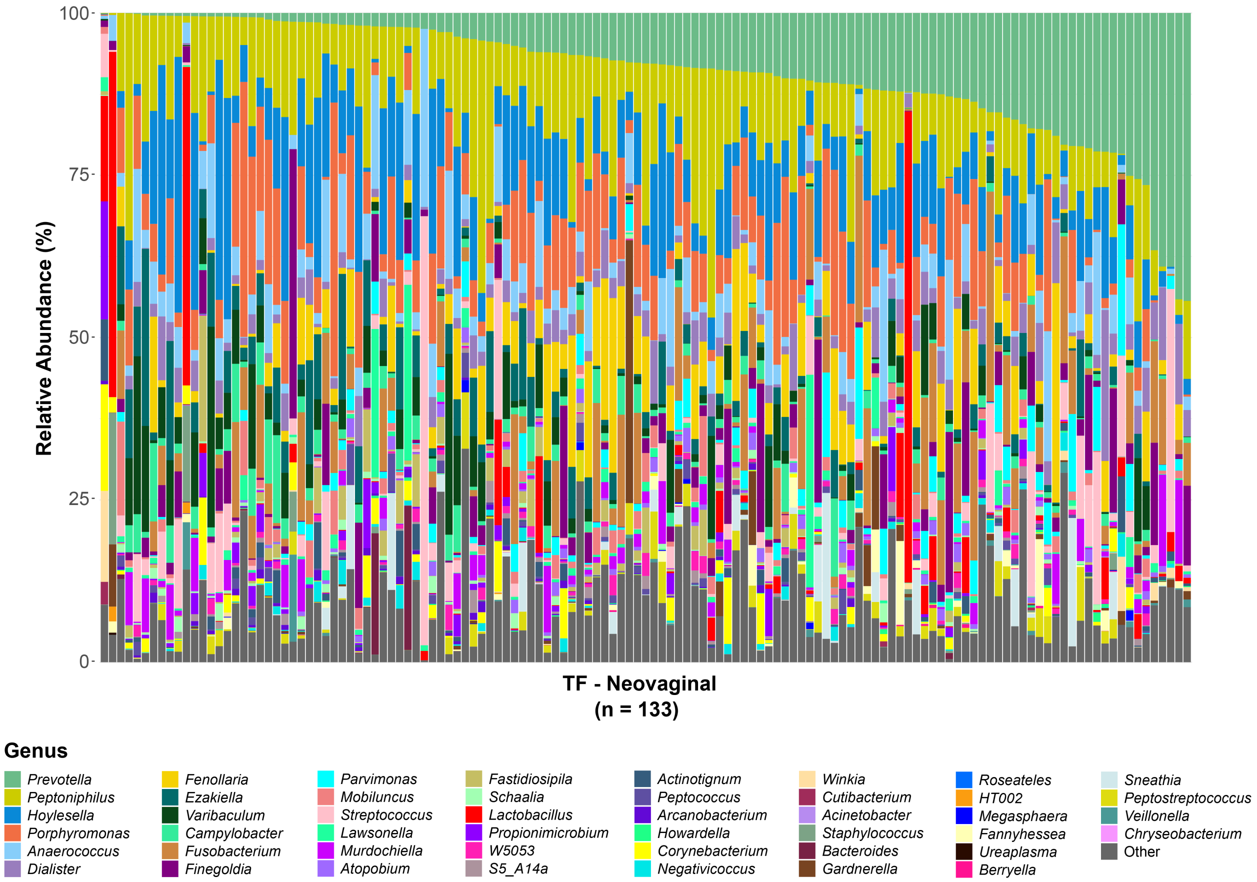


**Supplemental Figure 1. Relative abundances of bacteria in the TF neovagina.** Bacteria in genital swabs from transfeminine (TF, n=47 participants providing a total of n=133 visits) participants were characterized by 16S rRNA gene amplicon sequencing. The relative abundances of the top 30 most abundant genera in each anatomical location (by median relative abundance, identified in Figure 2) are displayed as stacked bar plots (total 46 genera). Each vertical bar represents one sample (n=133).

**Supplemental Table 3. Cytokine concentrations in neovaginal swab eluant.**

| | Median (pg/ml) | IQR (pg/ml) | Range (pg/ml) | LLOD (pg/ml) | LLOQ (pg/ml) | Samples detected |
|---|---|---|---|---|---|---|
| IL-1α | 3394 | 1,680 - 8,136 | 156.8 - 20,352 | | 4.80 | 100% |
| IL-8 | 451.1 | 83.5 - 1,565 | < 0.64 - 5,915 | | 0.64 | 99% |
| IL-1β | 33.1 | 4.80 - 823.6 | < 1.60 - 20,317 | 1.60 | 8.00 | 82% |
| MIP-1β | 5.75 | 2.25 - 6.80 | < 1.90 - 1,198 | 1.90 | 9.60 | 80% |
| IL-6 | 2.76 | < 0.64 - 15.5 | < 0.64 - 3,106 | | 0.64 | 73% |
| MIG | 12.5 | < 6.40 - 43.9 | < 6.40 - 1,857 | | 6.40 | 65% |
| RANTES | n/a | < 1.30 - 4.02 | < 1.30 - 858.0 | | 1.30 | 41% |
| TNFα | n/a | < 6.40 - 11.3 | < 6.40 - 446.7 | | 6.40 | 38% |
| IL-10 | n/a | < 2.60 - 3.42 | < 2.60 - 305.3 | | 2.60 | 28% |
| IFNγ | n/a | < 6.40 - 7.15 | < 6.40 - 31.8 | | 6.40 | 27% |
| MIP-1α | n/a | n/a | < 16.0 - 2,092 | | 16.0 | 19% |
| IL-22 | n/a | n/a | < 64.0 - 135.7 | | 64.0 | 18% |
| IP-10 | n/a | n/a | < 2.60 - 70.4 | 2.60 | 12.8 | 11% |
| IFN-α2 | n/a | n/a | < 39.6 - 70.1 | | 39.6 | 3% |
| IL-13 | n/a | n/a | < 31.9 | | 31.9 | 0% |
| IL-17A | n/a | n/a | < 34.3 - 75.4 | | 34.3 | 2% |
| IL-17E | n/a | n/a | < 180.2 | | 180.2 | 0% |
| IL-17F | n/a | n/a | < 31.7 - 117.1 | | 31.7 | 2% |
| GM-CSF | n/a | n/a | < 67.4 | | 67.4 | 0% |

IQR: Interquartile Range
LLOD: Lower Limit of Detection
LLOQ: Lower Limit of Quantification
n/a for median: >50% of samples < LLOD
n/a for IQR: >75% of samples < LLOD

### Supplemental Table 4. P-values associated with spearman correlation coefficients for correlations between relative abundances of the top 30 most abundant bacterial taxa in the neovagina (associated with Figure 3).

| | | *Dialister* | *Mobiluncus* | *Porphyromonas* | *Peptococcus* | *W5053* | *Fenollaria* | *Atopobium* | *Prevotella* | *Parvimonas* | *Fusobacterium* | *Howardella* | *Lawsonella* | *Streptococcus* | *Anaerococcus* | *Corynebacterium* | *Finegoldia* | *Schaalia* | *Lactobacillus* | *Arcanobacterium* | *Propionimicrobium* | *Negativicoccus* | *Campylobacter* | *Ezakiella* | *Actinotignum* | *Murdochiella* | *Hoylesella* | *Varibaculum* | *Peptoniphilus* | *S5-A14a* | *Fastidiosipila* |
|---|---|---|---|---|---|---|---|---|---|---|---|---|---|---|---|---|---|---|---|---|---|---|---|---|---|---|---|---|---|---|---|
| TC 1 | *Dialister* | 0 | 0.01 | 0.73 | 0.42 | 0.49 | 0.17 | 0.03 | 0.00 | 0.43 | 0.99 | 0.41 | 0.32 | 0.12 | 0.24 | 0.46 | 0.18 | 0.79 | 0.19 | 0.57 | 0.91 | 0.39 | 0.16 | 0.21 | 0.19 | 0.00 | 0.00 | 0.10 | 0.02 | 0.03 | 0.74 |
| | *Mobiluncus* | 0.01 | 0 | 0.00 | 0.00 | 0.00 | 0.00 | 0.02 | 0.44 | 0.80 | 0.75 | 0.26 | 0.02 | 0.00 | 0.00 | 0.01 | 0.00 | 0.89 | 0.02 | 0.37 | 0.08 | 0.67 | 0.04 | 0.02 | 0.69 | 0.10 | 0.19 | 0.02 | 0.01 | 0.00 | 0.00 |
| | *Porphyromonas* | 0.73 | 0.00 | 0 | 0.00 | 0.00 | 0.00 | 0.00 | 0.04 | 0.26 | 0.01 | 0.17 | 0.05 | 0.00 | 0.03 | 0.04 | 0.00 | 0.22 | 0.00 | 0.01 | 0.11 | 0.34 | 0.00 | 0.00 | 0.57 | 0.94 | 0.25 | 0.24 | 0.06 | 0.00 | 0.00 |
| | *Peptococcus* | 0.42 | 0.00 | 0.00 | 0 | 0.00 | 0.00 | 0.00 | 0.76 | 0.02 | 0.02 | 0.91 | 0.12 | 0.00 | 0.19 | 0.03 | 0.00 | 0.97 | 0.01 | 0.00 | 0.56 | 0.99 | 0.05 | 0.08 | 0.43 | 0.84 | 0.64 | 0.57 | 0.48 | 0.00 | 0.00 |
| | *W5053* | 0.49 | 0.00 | 0.00 | 0.00 | 0 | 0.00 | 0.00 | 0.38 | 0.68 | 0.42 | 0.34 | 0.28 | 0.00 | 0.16 | 0.07 | 0.00 | 0.56 | 0.02 | 0.00 | 0.60 | 0.37 | 0.04 | 0.02 | 0.13 | 0.12 | 0.64 | 0.58 | 0.03 | 0.00 | 0.00 |
| | *Fenollaria* | 0.17 | 0.00 | 0.00 | 0.00 | 0.00 | 0 | 0.00 | 0.49 | 0.00 | 0.00 | 0.48 | 0.56 | 0.00 | 0.02 | 0.00 | 0.00 | 0.20 | 0.05 | 0.35 | 0.44 | 0.59 | 0.03 | 0.56 | 0.45 | 0.03 | 0.24 | 0.20 | 0.58 | 0.03 | 0.01 |
| | *Atopobium* | 0.03 | 0.02 | 0.00 | 0.00 | 0.00 | 0.00 | 0 | 0.02 | 0.00 | 0.00 | 0.01 | 0.12 | 0.05 | 0.01 | 0.01 | 0.07 | 0.92 | 0.58 | 0.37 | 0.78 | 0.17 | 0.61 | 0.35 | 0.04 | 0.72 | 0.01 | 0.20 | 0.46 | 0.29 | 0.53 |
| TC 2 | *Prevotella* | 0.00 | 0.44 | 0.04 | 0.76 | 0.38 | 0.49 | 0.02 | 0 | 0.00 | 0.02 | 0.00 | 0.46 | 0.91 | 0.02 | 0.01 | 0.76 | 0.00 | 0.01 | 0.00 | 0.00 | 0.40 | 0.00 | 0.00 | 0.44 | 0.97 | 0.14 | 0.00 | 0.04 | 0.10 | 0.02 |
| | *Parvimonas* | 0.43 | 0.80 | 0.26 | 0.02 | 0.68 | 0.00 | 0.00 | 0.00 | 0 | 0.00 | 0.00 | 0.00 | 0.98 | 0.78 | 0.00 | 0.00 | 0.27 | 0.60 | 0.03 | 0.08 | 0.06 | 0.02 | 0.04 | 0.85 | 0.02 | 0.99 | 0.00 | 0.00 | 0.19 | 0.12 |
| | *Fusobacterium* | 0.99 | 0.75 | 0.01 | 0.02 | 0.42 | 0.00 | 0.00 | 0.02 | 0.00 | 0 | 0.00 | 0.04 | 0.45 | 0.00 | 0.00 | 0.00 | 0.58 | 0.48 | 0.02 | 0.05 | 0.11 | 0.69 | 0.09 | 0.12 | 0.42 | 0.31 | 0.00 | 0.00 | 0.35 | 0.11 |
| | *Howardella* | 0.41 | 0.26 | 0.17 | 0.91 | 0.34 | 0.48 | 0.01 | 0.00 | 0.00 | 0.00 | 0 | 0.91 | 0.00 | 0.18 | 0.58 | 0.28 | 0.01 | 0.01 | 0.04 | 0.19 | 0.30 | 0.04 | 0.00 | 0.64 | 0.06 | 0.42 | 0.03 | 0.00 | 0.01 | 0.04 |
| | *Lawsonella* | 0.32 | 0.02 | 0.05 | 0.12 | 0.28 | 0.56 | 0.12 | 0.46 | 0.00 | 0.04 | 0.91 | 0 | 0.17 | 0.06 | 0.12 | 0.80 | 0.14 | 0.01 | 0.06 | 0.24 | 0.00 | 0.00 | 0.14 | 0.00 | 0.57 | 0.69 | 0.12 | 0.30 | 0.86 | 0.26 |
| TC 3 | *Streptococcus* | 0.12 | 0.00 | 0.00 | 0.00 | 0.00 | 0.00 | 0.05 | 0.91 | 0.98 | 0.45 | 0.00 | 0.17 | 0 | 0.00 | 0.00 | 0.00 | 0.02 | 0.09 | 0.42 | 0.73 | 0.80 | 0.08 | 0.05 | 0.10 | 0.06 | 0.69 | 0.44 | 0.05 | 0.14 | 0.00 |
| | *Anaerococcus* | 0.24 | 0.00 | 0.03 | 0.19 | 0.16 | 0.02 | 0.01 | 0.02 | 0.78 | 0.00 | 0.18 | 0.06 | 0.00 | 0 | 0.00 | 0.00 | 0.00 | 0.16 | 0.03 | 0.00 | 0.40 | 0.58 | 0.91 | 0.78 | 0.13 | 0.56 | 0.00 | 0.69 | 0.71 | 0.28 |
| | *Corynebacterium* | 0.46 | 0.01 | 0.04 | 0.03 | 0.07 | 0.00 | 0.01 | 0.01 | 0.00 | 0.00 | 0.58 | 0.12 | 0.00 | 0.00 | 0 | 0.00 | 0.02 | 0.10 | 0.02 | 0.00 | 0.01 | 0.91 | 0.46 | 0.41 | 0.35 | 0.47 | 0.01 | 0.16 | 0.18 | 0.95 |
| | *Finegoldia* | 0.18 | 0.00 | 0.00 | 0.00 | 0.00 | 0.00 | 0.07 | 0.76 | 0.00 | 0.00 | 0.28 | 0.80 | 0.00 | 0.00 | 0.00 | 0 | 0.10 | 0.07 | 0.84 | 0.58 | 0.64 | 0.02 | 0.01 | 0.48 | 0.83 | 0.92 | 0.22 | 0.04 | 0.20 | 0.05 |
| | *Schaalia* | 0.79 | 0.89 | 0.22 | 0.97 | 0.56 | 0.20 | 0.92 | 0.00 | 0.27 | 0.58 | 0.01 | 0.14 | 0.02 | 0.00 | 0.02 | 0.10 | 0 | 0.11 | 0.10 | 0.02 | 0.39 | 0.44 | 0.72 | 0.38 | 0.57 | 0.00 | 0.00 | 0.18 | 0.83 | 0.52 |
| | *Lactobacillus* | 0.19 | 0.02 | 0.00 | 0.01 | 0.02 | 0.05 | 0.58 | 0.01 | 0.60 | 0.48 | 0.01 | 0.01 | 0.09 | 0.16 | 0.10 | 0.07 | 0.11 | 0 | 0.09 | 0.68 | 0.75 | 0.00 | 0.00 | 0.09 | 0.05 | 0.06 | 0.37 | 0.00 | 0.07 | 0.19 |
| TC 4 | *Arcanobacterium* | 0.57 | 0.37 | 0.01 | 0.00 | 0.00 | 0.35 | 0.37 | 0.00 | 0.03 | 0.02 | 0.04 | 0.06 | 0.42 | 0.03 | 0.02 | 0.84 | 0.10 | 0.09 | 0 | 0.00 | 0.00 | 0.00 | 0.00 | 0.01 | 0.31 | 0.26 | 0.00 | 0.00 | 0.00 | 0.00 |
| | *Propionimicrobium* | 0.91 | 0.08 | 0.11 | 0.56 | 0.60 | 0.44 | 0.78 | 0.00 | 0.08 | 0.05 | 0.19 | 0.24 | 0.73 | 0.00 | 0.00 | 0.58 | 0.02 | 0.68 | 0.00 | 0 | 0.00 | 0.00 | 0.00 | 0.01 | 0.35 | 0.24 | 0.00 | 0.00 | 0.00 | 0.00 |
| | *Negativicoccus* | 0.39 | 0.67 | 0.34 | 0.99 | 0.37 | 0.59 | 0.17 | 0.40 | 0.06 | 0.11 | 0.30 | 0.00 | 0.80 | 0.40 | 0.01 | 0.64 | 0.39 | 0.75 | 0.00 | 0.00 | 0 | 0.00 | 0.00 | 0.00 | 0.26 | 0.85 | 0.00 | 0.19 | 0.01 | 0.00 |
| | *Campylobacter* | 0.16 | 0.04 | 0.00 | 0.05 | 0.04 | 0.03 | 0.61 | 0.00 | 0.02 | 0.69 | 0.04 | 0.00 | 0.08 | 0.58 | 0.91 | 0.02 | 0.44 | 0.00 | 0.00 | 0.00 | 0.00 | 0 | 0.00 | 0.01 | 0.01 | 0.00 | 0.00 | 0.00 | 0.00 | 0.00 |
| | *Ezakiella* | 0.21 | 0.02 | 0.00 | 0.08 | 0.02 | 0.56 | 0.35 | 0.00 | 0.04 | 0.09 | 0.00 | 0.14 | 0.05 | 0.91 | 0.46 | 0.01 | 0.72 | 0.00 | 0.00 | 0.00 | 0.00 | 0.00 | 0 | 0.19 | 0.00 | 0.16 | 0.00 | 0.00 | 0.00 | 0.00 |
| | *Actinotignum* | 0.19 | 0.69 | 0.57 | 0.43 | 0.13 | 0.45 | 0.04 | 0.44 | 0.85 | 0.12 | 0.64 | 0.00 | 0.10 | 0.78 | 0.41 | 0.48 | 0.38 | 0.09 | 0.01 | 0.01 | 0.00 | 0.01 | 0.19 | 0 | 0.89 | 0.88 | 0.05 | 0.55 | 0.42 | 0.04 |
| | *Murdochiella* | 0.00 | 0.10 | 0.94 | 0.84 | 0.12 | 0.03 | 0.72 | 0.97 | 0.02 | 0.42 | 0.06 | 0.57 | 0.06 | 0.13 | 0.35 | 0.83 | 0.57 | 0.05 | 0.31 | 0.35 | 0.26 | 0.01 | 0.00 | 0.89 | 0 | 0.00 | 0.00 | 0.00 | 0.00 | 0.01 |
| | *Hoylesella* | 0.00 | 0.19 | 0.25 | 0.64 | 0.64 | 0.24 | 0.01 | 0.14 | 0.99 | 0.31 | 0.42 | 0.69 | 0.69 | 0.56 | 0.47 | 0.92 | 0.00 | 0.06 | 0.26 | 0.24 | 0.85 | 0.00 | 0.16 | 0.88 | 0.00 | 0 | 0.00 | 0.19 | 0.07 | 0.72 |
| | *Varibaculum* | 0.10 | 0.02 | 0.24 | 0.57 | 0.58 | 0.20 | 0.20 | 0.00 | 0.00 | 0.00 | 0.03 | 0.12 | 0.44 | 0.00 | 0.01 | 0.22 | 0.00 | 0.37 | 0.00 | 0.00 | 0.00 | 0.00 | 0.00 | 0.05 | 0.00 | 0.00 | 0 | 0.00 | 0.00 | 0.01 |
| | *Peptoniphilus* | 0.02 | 0.01 | 0.06 | 0.48 | 0.03 | 0.58 | 0.46 | 0.04 | 0.00 | 0.00 | 0.00 | 0.30 | 0.05 | 0.69 | 0.16 | 0.04 | 0.18 | 0.00 | 0.00 | 0.00 | 0.19 | 0.00 | 0.00 | 0.55 | 0.00 | 0.19 | 0.00 | 0 | 0.00 | 0.02 |
| | *S5-A14a* | 0.03 | 0.00 | 0.00 | 0.00 | 0.00 | 0.03 | 0.29 | 0.10 | 0.19 | 0.35 | 0.01 | 0.86 | 0.14 | 0.71 | 0.18 | 0.20 | 0.83 | 0.07 | 0.00 | 0.00 | 0.01 | 0.00 | 0.00 | 0.42 | 0.00 | 0.07 | 0.00 | 0.00 | 0 | 0.00 |
| | *Fastidiosipila* | 0.74 | 0.00 | 0.00 | 0.00 | 0.00 | 0.01 | 0.53 | 0.02 | 0.12 | 0.11 | 0.04 | 0.26 | 0.00 | 0.28 | 0.95 | 0.05 | 0.52 | 0.19 | 0.00 | 0.00 | 0.00 | 0.00 | 0.00 | 0.04 | 0.01 | 0.72 | 0.01 | 0.02 | 0.00 | 0 |

**Supplemental Table 5. P-values associated with spearman correlation coefficients for correlations between bacterial taxa relative abundance and cytokine concentration, for the top 30 most abundant bacterial taxa in the neovagina (associated with Figure 3).**

| | | IL-1α | IL-1β | IL-6 | IL-8 | MIG | MIP-1β | RANTES |
|---|---|---|---|---|---|---|---|---|
| TC 1 | *Dialister* | 0.487 | 0.523 | 0.340 | 0.783 | 0.505 | 0.959 | 0.886 |
| | *Mobiluncus* | 0.535 | 0.646 | 0.059 | 0.106 | **0.021** | 0.524 | 0.270 |
| | *Porphyromonas* | 0.130 | **0.000** | 0.374 | 0.732 | 0.246 | 0.230 | 0.676 |
| | *Peptococcus* | 0.176 | **0.013** | 0.430 | 0.730 | 0.091 | 0.651 | 0.737 |
| | *W5053* | 0.697 | 0.855 | 0.422 | 0.073 | **0.008** | 0.262 | 0.179 |
| | *Fenollaria* | 0.977 | 0.750 | 0.093 | 0.252 | 0.288 | 0.651 | 0.155 |
| | *Atopobium* | 0.082 | 0.054 | 0.953 | 0.885 | 0.885 | 0.328 | 0.712 |
| TC 2 | *Prevotella* | **0.000** | 0.473 | 0.551 | 0.236 | **0.039** | 0.279 | 0.473 |
| | *Parvimonas* | **0.004** | **0.000** | **0.019** | **0.013** | 0.268 | **0.029** | 0.059 |
| | *Fusobacterium* | **0.000** | **0.021** | **0.046** | **0.032** | **0.035** | **0.026** | 0.063 |
| | *Howardella* | **0.013** | 0.224 | 0.083 | **0.028** | **0.029** | **0.047** | **0.019** |
| | *Lawsonella* | **0.019** | **0.000** | **0.026** | **0.004** | 0.378 | **0.002** | **0.008** |
| TC 3 | *Streptococcus* | 0.917 | 0.844 | **0.000** | **0.003** | **0.000** | **0.006** | 0.055 |
| | *Anaerococcus* | 0.486 | 0.487 | **0.022** | 0.054 | **0.040** | **0.042** | 0.340 |
| | *Corynebacterium* | 0.290 | 0.524 | 0.245 | 0.455 | 0.116 | 0.473 | 0.289 |
| | *Finegoldia* | 0.821 | 0.651 | 0.266 | 0.183 | **0.003** | 0.146 | 0.263 |
| | *Schaalia* | 0.665 | **0.019** | **0.008** | **0.004** | 0.068 | **0.006** | 0.051 |
| | *Lactobacillus* | 0.285 | **0.032** | 0.267 | 0.710 | 0.267 | 0.746 | 0.455 |
| TC 4 | *Arcanobacterium* | 0.059 | 0.700 | 0.737 | 0.151 | 0.047 | 0.651 | 0.279 |
| | *Propionimicrobium* | 0.035 | 0.651 | 0.791 | 0.177 | 0.291 | 0.932 | 0.243 |
| | *Negativicoccus* | 0.245 | 0.032 | 0.288 | 0.008 | 0.122 | 0.021 | 0.025 |
| | *Campylobacter* | 0.487 | 0.220 | 0.524 | 0.059 | 0.328 | 0.470 | 0.487 |
| | *Ezakiella* | 0.032 | 0.554 | 0.262 | 0.013 | 0.000 | 0.041 | 0.003 |
| | *Actinotignum* | 0.551 | 0.262 | 0.710 | 0.288 | 0.550 | 0.800 | 0.855 |
| | *Murdochiella* | 0.374 | 0.142 | 0.008 | 0.019 | 0.105 | 0.032 | 0.003 |
| | *Hoylesella* | 0.971 | 0.855 | 0.607 | 0.913 | 0.651 | 0.743 | 0.950 |
| | *Varibaculum* | 0.029 | 0.539 | 0.797 | 0.094 | 0.712 | 0.710 | 0.374 |
| | *Peptoniphilus* | 0.047 | 0.719 | 0.059 | 0.029 | 0.019 | 0.132 | 0.094 |
| | *S5-A14a* | 0.285 | 0.901 | 0.370 | 0.081 | 0.059 | 0.486 | 0.075 |
| | *Fastidiosipila* | 0.051 | 0.166 | 0.014 | 0.001 | 0.003 | 0.018 | 0.018 |

**Supplemental Table 6. Median abundance of each Taxa Cluster (TC), by co-variate.**

| | | Total median abundance (%) | | | |
|---|---|---|---|---|---|
| | | TC1 | TC2 | TC3 | TC4 |
| Self-reported neovaginal symptoms, previous 7 days | None | 16.50 | 13.49 | 7.96 | 32.29 |
| | Bleeding | 21.38 | 26.77 | 9.52 | 26.39 |
| | Discharge | 8.95 | 22.02 | 30.02 | 24.27 |
| | Itch | 18.57 | 14.94 | 6.12 | 30.32 |
| | Malodour | 19.02 | 6.35 | 9.16 | 50.67 |
| | Pain | 6.81 | 12.42 | 28.31 | 45.00 |
| Pre-vaginoplasty circ. status | Uncircumcised | 21.96 | 15.83 | 5.93 | 26.93 |
| | Circumcised | 14.54 | 9.98 | 10.99 | 39.50 |
| Neovaginal behavioural type | Minimal exposures | 21.56 | 9.35 | 4.80 | 39.48 |
| | Dilating only | 17.04 | 12.58 | 9.17 | 33.27 |
| | Douching & dilating | 18.56 | 14.25 | 5.20 | 34.81 |
| | Diverse exposures | 9.18 | 18.62 | 17.53 | 25.10 |

**Supplemental Table 7. Beta coefficient and p-values of Zero-inflated beta regression model with random effects to account for repeated measurements, by co-variate.**

| | | **TC1** | | **TC2** | | **TC3** | | **TC4** | |
|---|---|---|---|---|---|---|---|---|---|
| | | Beta Coefficient | p-value | Beta Coefficient | p-value | Beta Coefficient | p-value | Beta Coefficient | p-value |
| | Age | 0.004 | 0.471 | -0.005 | 0.286 | -0.002 | 0.790 | -0.002 | 0.581 |
| | Time since vaginoplasty | -0.014 | 0.333 | 0.011 | 0.340 | 0.009 | 0.585 | -0.012 | 0.307 |
| | pH | 0.132 | 0.063 | -0.026 | 0.661 | -0.143 | 0.920 | 0.134 | **0.026** |
| *vs. uncircumcised* | Circumcised | -0.415 | **< 0.001** | -0.492 | **< 0.001** | 0.389 | **< 0.001** | 0.447 | **< 0.001** |
| *vs. minimal exposures* | Dilating only | -0.141 | 0.394 | 0.391 | **0.014** | 0.289 | 0.209 | -0.265 | 0.072 |
| | Douching & dilating | -0.391 | 0.384 | 0.491 | **0.003** | 0.009 | 0.97 | -0.306 | **0.05** |
| | Diverse exposures | -0.866 | **< 0.001** | 0.683 | **< 0.001** | 0.88 | **< 0.001** | -0.634 | **< 0.001** |
| *vs. no reported symptoms* | Bleeding | -0.075 | 0.704 | 0.254 | 0.076 | 0.099 | 0.686 | -0.041 | 0.759 |
| | Discharge | -0.15 | 0.567 | -0.016 | 0.93 | 0.058 | 0.829 | 0.077 | 0.705 |
| | Itch | -0.012 | 0.959 | -0.056 | 0.785 | 0.242 | 0.438 | 0.046 | 0.824 |
| | Malodour | -0.04 | 0.79 | -0.434 | **0.005** | 0.002 | 0.992 | 0.464 | **< 0.001** |
| | Pain | -0.313 | 0.561 | -0.403 | 0.296 | 0.052 | 0.916 | 1.152 | **< 0.001** |

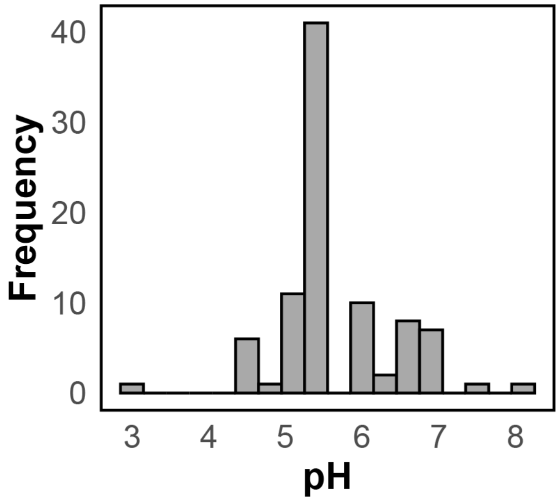


**Supplemental Figure 2: Distribution of self-reported neovaginal pH of n=94 samples collected by n=47 TF participants.** Participants self-assessed pH by rolling a neovaginal swab onto a colorimetric pH strip (provided).

**Supplemental Table 8. Prevalence of self-reported neovaginal symptoms in the past 7 days, at study visits.**

| | Visits (n=133) |
|---|---|
| **Symptom** (n, %) | |
| No symptoms | 98 (73.68) |
| Bleeding | 10 (7.52) |
| Discharge | 7 (5.26) |
| Itch/Burn | 6 (4.51) |
| Malodour | 19 (14.29) |
| Pain | 2 (1.50) |
